# SNX-BAR proteins 5 and 6 are required for NCOA7-AS antiviral activity against influenza A virus

**DOI:** 10.1101/2025.04.01.646557

**Authors:** Mary Arnaud-Arnould, Antoine Rebendenne, Marine Tauziet, Serge Urbach, Khadija El Koulali, Emiliano P. Ricci, Mélanie Wencker, Olivier Moncorgé, Mickaël Blaise, Caroline Goujon

**Author notes:** Contributed equally to the work (Mary Arnaud-Arnould, Antoine Rebendenne).

## Abstract

Interferon-induced antiviral proteins act on the first line of defence against viruses, including influenza A virus (IAV). Among those, the short isoform of Nuclear Receptor Coactivator 7 (NCOA7-AS) has been shown to inhibit IAV entry and more especially the fusion between endosomal and viral membranes. Ectopic expression of NCOA7-AS leads to the increased acidification of the endolysosomal pathway, putatively through NCOA7-AS interaction with the vacuolar ATPase (V-ATPase). However, the precise mechanism of action of NCOA7-AS is not fully understood. Here, we identified the cellular partners of NCOA7-AS by mass spectrometry, including the V-ATPase subunits known to interact with NCOA7-AS. A point mutation disrupting the interaction with the V-ATPase led to loss of antiviral activity against IAV, demonstrating that V-ATPase engagement is required for the antiviral phenotype. Moreover, sorting nexin (SNX) 1/2/5/6 were identified as novel partners of NCOA7-AS. These proteins are involved in retrograde transport of cellular cargoes from endosomes to the trans-Golgi network. We revealed that SNX5/6 interacted directly with NCOA7-AS and were essential for NCOA7-AS antiviral activity against IAV. Interestingly, crystal structures of NCOA7-AS/SNX5 complexes showed that the SNX5-interaction motif in NCOA7-AS was similar to those found in known cargoes of SNX5/6. In addition, we could pinpoint several critical residues that were important for binding and antiviral activity. Collectively, our study identifies novel essential partners for NCOA7-AS antiviral activity and the structural basis for their interaction.

## Introduction

Seasonal influenza A viruses (IAV) pose a major threat to global health, putting significant pressure on healthcare systems and causing significant morbidity and mortality each year, according to the World Health Organization. IAV are respiratory viruses, belonging to the *Orthomyxoviridae* family, that cause mild to moderate upper respiratory tracts illnesses. Successful infection of host cells by IAV depends on a complex, multi-step entry process. The hemagglutinin (HA) protein of human-adapted strains of IAV interacts preferentially with glycoconjugates bearing terminal α-2,6-linked sialic acids (reviewed in (Dou et al., 2018)). Following this, IAV particle is internalized through endocytosis (Matlin et al., 1981). This early step requires numerous host factors involved in the endocytic pathway, as well as the vacuolar ATPase (V-ATPase), which drives the progressive acidification of the endosomal lumen. The V-ATPase is a multi-subunit complex, composed of two distinct domains: the membrane-associated V0 domain, which is responsible for proton translocation to the lumen of the vesicle, and the cytoplasmic V1 domain, which provides energy by ATP hydrolysis (reviewed in (Cotter et al., 2015)). Endosomal acidification by the V-ATPase is critical for HA acidification, fusion peptide exposure and subsequent HA-mediated fusion between viral and endosomal membranes. Furthermore, luminal acidification activates the viral M2 ion channel, leading to proton influx and acidification within the viral particle (Pinto et al., 1992; Wharton et al., 1994; Pinto and Lamb, 2006; Stauffer et al., 2014). This induces conformational changes in the matrix protein M1, disrupting the viral capsid and the interaction between M1 and the viral ribonucleoproteins complexes (vRNPs), leading to the subsequent release of vRNPs in the cytoplasm (Zhirnov, 1990; Helenius, 1992).

IAV entry is a critical step targeted by many potent antiviral proteins, including some coded by interferon-stimulated genes (ISGs) that are part of the interferon (IFN) system. The IFNs act as one of the first lines of defence against viral infections (reviewed in (Villalón-Letelier et al., 2017; McKellar et al., 2021)). Notably, the 219-amino acid, interferon-inducible, short isoform of Nuclear Receptor Coactivator 7 (NCOA7), termed NCOA7-Alternative-Start (NCOA7-AS; also referred to as NCOA7 isoform 4 or NCOA7-B; (Yu et al., 2015)), has been shown to inhibit IAV entry (Doyle et al., 2018) as well as endosome-mediated coronavirus entry (Khan et al., 2021; Wickenhagen et al., 2021). In contrast to this interferon-inducible, antiviral isoform, the longer NCOA7 isoform (herein referred to as NCOA7-FL) is constitutively expressed and has been primarily implicated in neuronal and lysosomal homeostasis (Merkulova et al., 2015; Doyle et al., 2018; Castroflorio et al., 2021). NCOA7-FL and NCOA7-AS belong to the Tre2/Bub2/Cdc16 (TBC), lysin motif (LysM), domain catalytic (TLDc)-containing family of proteins. Despite their name, no enzymatic activity has ever been reported for TLDc-bearing proteins (Blaise et al., 2012), but they have been shown to protect cells from oxidative stress, by an unknown mechanism (Finelli and Oliver, 2017). Interestingly, the TLDc proteins are known interactors of the V-ATPase (Merkulova et al., 2015; Eaton et al., 2021). Ectopic expression of NCOA7-AS increases acidification of the endolysosomal pathway, supposedly through NCOA7-AS interaction with the V-ATPase (Doyle et al., 2018). NCOA7-AS ectopic expression is also associated with a decrease in IAV/endolysosomal membrane fusion (Doyle et al., 2018). However, the precise molecular mechanism of action of NCOA7-AS is yet to be unraveled.

Here, we employed mass spectrometry to identify the cellular partners of NCOA7-AS. This approach confirmed NCOA7-AS’ interaction with multiple V-ATPase subunits. Disruption of this interaction by a single amino acid substitution abolished antiviral activity against IAV, underscoring the importance of V-ATPase engagement in mediating the antiviral effect. Additionally, sorting nexins SNX1, SNX2, SNX5, and SNX6 were identified as novel binding partners of NCOA7-AS. These proteins are known to play roles in retrograde transport of cellular cargoes to the trans-Golgi network (Burd and Cullen, 2014). We further demonstrated that SNX5 and SNX6 bind directly to NCOA7-AS’ amino-terminal domain and were required for IAV inhibition and the overacidification of the endolysosomal compartment. Moreover, crystal structures of NCOA7-AS in complex with SNX5 revealed that the SNX5-binding motif in NCOA7-AS resembles those found in established SNX5/6 cargoes (Simonetti et al., 2019). Collectively, these findings strongly suggest that NCOA7-AS uses SNX proteins to suppress IAV replication.

## Results

### The interaction between NCOA7-AS and V1 subunits of the V-ATPase is required for antiviral activity

TLDc proteins are known interactors of the V-ATPase (Merkulova et al., 2015; Eaton et al., 2021). Notably, both NCOA7-FL and NCOA7-AS, which share a C-terminal TLDc domain, have been shown to interact with V1 subunits of the V-ATPase (ATP6V1B1/2 and also A, C and G) and to promote endolysosomal acidification in cells (Merkulova et al., 2015; Doyle et al., 2018; Castroflorio et al., 2021). As previously shown, several residues within TLDc proteins are highly conserved across species and are critical for V-ATPase binding (Eaton et al., 2021). Notably, mutations of glycine 815, glycine 845 and glycine 896 in mouse NCOA7-FL abolished interaction with the V-ATPase (Eaton et al., 2021). Among these 3 residues, only glycine 815 is exposed on the surface of the TLDc, therefore the focus was made on the equivalent glycine in NCOA7-AS (glycine 91). As shown by docking of NCOA7-AS AlphaFold model on the Cryo-EM structure of human V-ATPase bound to another TLDc domain containing protein, mEAK7 (Wang et al., 2022), this glycine 91 is present on a short loop in close proximity to the V-ATPase subunit B (Figure 1A). In order to study its importance, glycine 91 was mutated to an alanine (G91A). Lung-derived A549 cells were transduced to stably express a negative control (Firefly), wild-type NCOA7-AS (NCOA7-AS-WT) or NCOA7-AS-G91A mutant. Co-immunoprecipitation experiments confirmed that NCOA7-AS interacted with V1 subunits of the V-ATPase, namely ATP6V1A, ATP6V1B2 and ATP6V1E1, while NCOA7-AS-G91A mutant was unable to interact with these subunits (Figure 1B).

**Figure 1:**
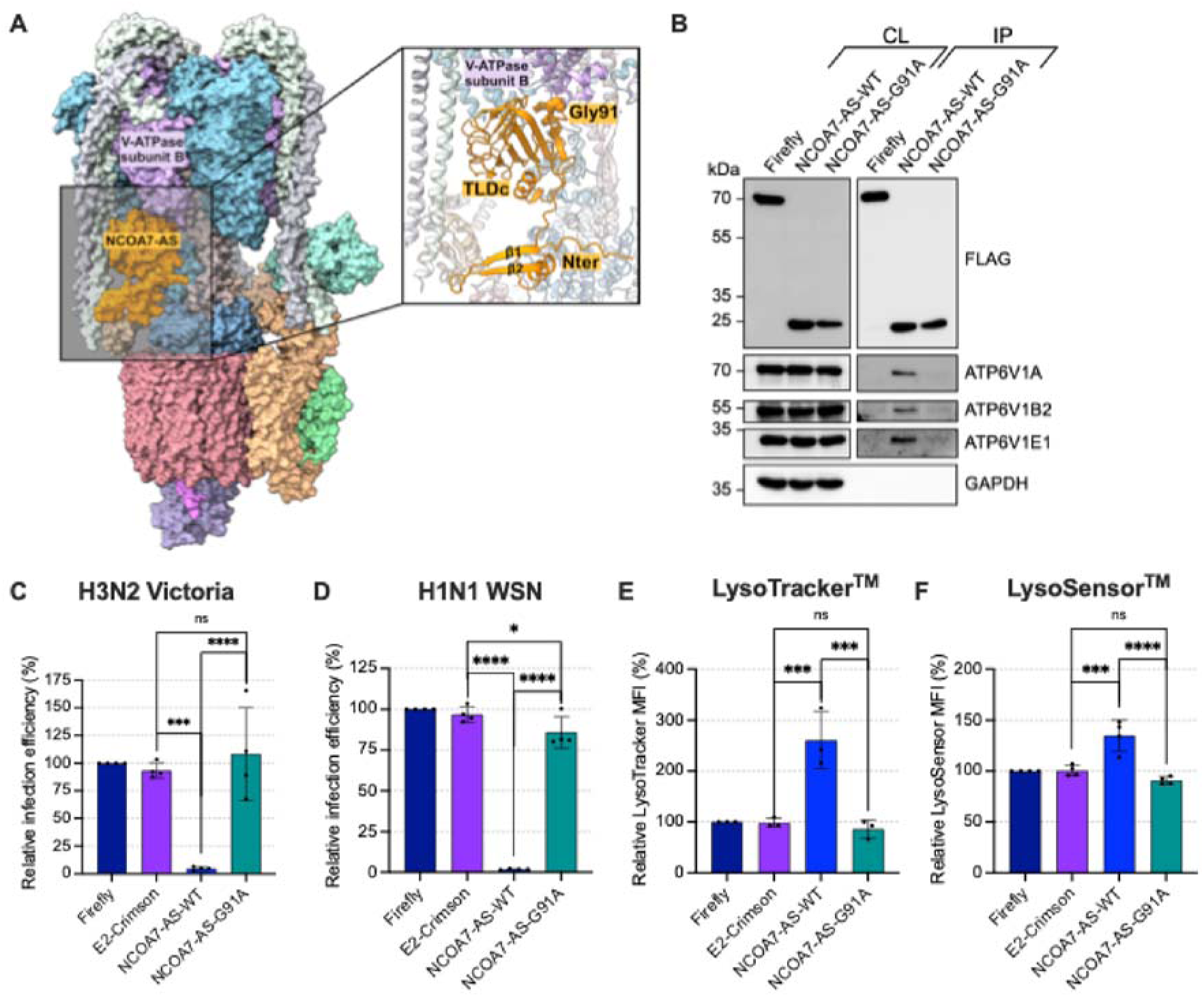
NCOA7-AS G91 is critical for V-ATPase binding, antiviral activity against IAV, and endolysosomal overacidification. **A.** Docking of NCOA7-AS AlphaFold model on the Cryo-EM structure of human V-ATPase bound to mEAK7, a TLDc domain containing protein (PDB:7unf). The NCOA7-AS model (in orange) was superposed onto the TLDc domain of mEAK7 using ChimeraX. The zoom panel indicates that the loop harboring NCOA7-AS G91, shown as spheres, is in close proximity to the V-ATPase B subunit. **B-F.** A549 cells were transduced with lentiviral vectors to stably express FLAG-tagged Firefly (a negative control), -E2-crimson (another negative control), -NCOA7-AS-WT or - NCOA7-AS-G91A. **B.** Cells were lysed and the FLAG-tagged proteins immunoprecipitated and subsequently analyzed by immunoblotting using the indicated antibodies; GAPDH served as a loading control (a representative immunoblot is shown out of 3 independent experiments). **C-D.** Cells were infected with IAV strains encoding a Nanoluciferase reporter gene: either A/Victoria/3/75 at MOI 0.5 (**C**) or A/WSN/1933 at MOI 0.25 (**D**). The relative infection efficiency was measured by monitoring Nanoluciferase activity 8h post-infection. **E-F.** Cells were incubated with LysoTracker^TM^ Green DND-26 (**E**) or with LysoSensor^TM^ Green DND-189 (**F**) for 1h at 37°C and the mean fluorescence intensity (MFI) was subsequently analyzed by flow cytometry. (**C-F**) Results were normalized to 100% for the control condition (Firefly), and the mean and SD of three independent experiments are shown; one-way ANOVA.

To determine whether interactions of NCOA7-AS with the V-ATPase were indeed required for antiviral activity and overacidification of the endolysosomal compartment, NCOA7-AS-G91A antiviral activity was tested against Nanoluciferase-encoding influenza reporter viruses A/Victoria/3/75 (H3N2) and A/WSN/1933 (H1N1) in single-cycle experiments. Ectopic expression of NCOA7-AS-WT led to a strong inhibition of A/Victoria/3/75 and A/WSN/1933 infection (Figure 1C-D), while NCOA7-AS-G91A mutant showed almost no antiviral activity. Furthermore, the endosomal/lysosomal compartment size and relative acidification levels were assessed using LysoTracker^TM^ and LysoSensor^TM^ reagents, respectively. This confirmed that NCOA7-AS-WT induced a significant expansion of the acidic compartments (Figure 1E) and a decrease in endosomal/lysosomal pH (Figure 1F). In contrast, NCOA7-AS-G91A mutant had no impact as the values were similar to the negative control conditions, strongly arguing that the interaction of NCOA7-AS with the V-ATPase is required for antiviral activity against IAV and overacidification of the endolysosomal compartment.

### SNX5 and SNX6 are cellular partners of NCOA7-AS required for its function

To identify additional binding partners of NCOA7-AS and gain further insight into its mode of action, 3xFLAG-tagged NCOA7-AS was stably expressed in U87-MG glioblastoma cells, which are highly permissive to lentiviral transduction and support high levels of transgene expression. The protein was then immunoprecipitated, and the resulting complexes were analyzed by mass spectrometry (Supplementary Table 1). The interaction between NCOA7-AS and several V1 subunits of the V-ATPase, namely ATP6V1A, ATP6V1B2, ATP6V1C1, ATP6V1D, ATP6V1E1, and ATP6V1G1, was confirmed (Figure 2A; Supplementary Table 1). The NCOA7-AS interactome additionally contained four members of the sorting nexins (SNX) family, SNX1, SNX2, SNX5 and SNX6, which were particularly enriched (Figure 2A; Supplementary Table 1).

**Figure 2:**
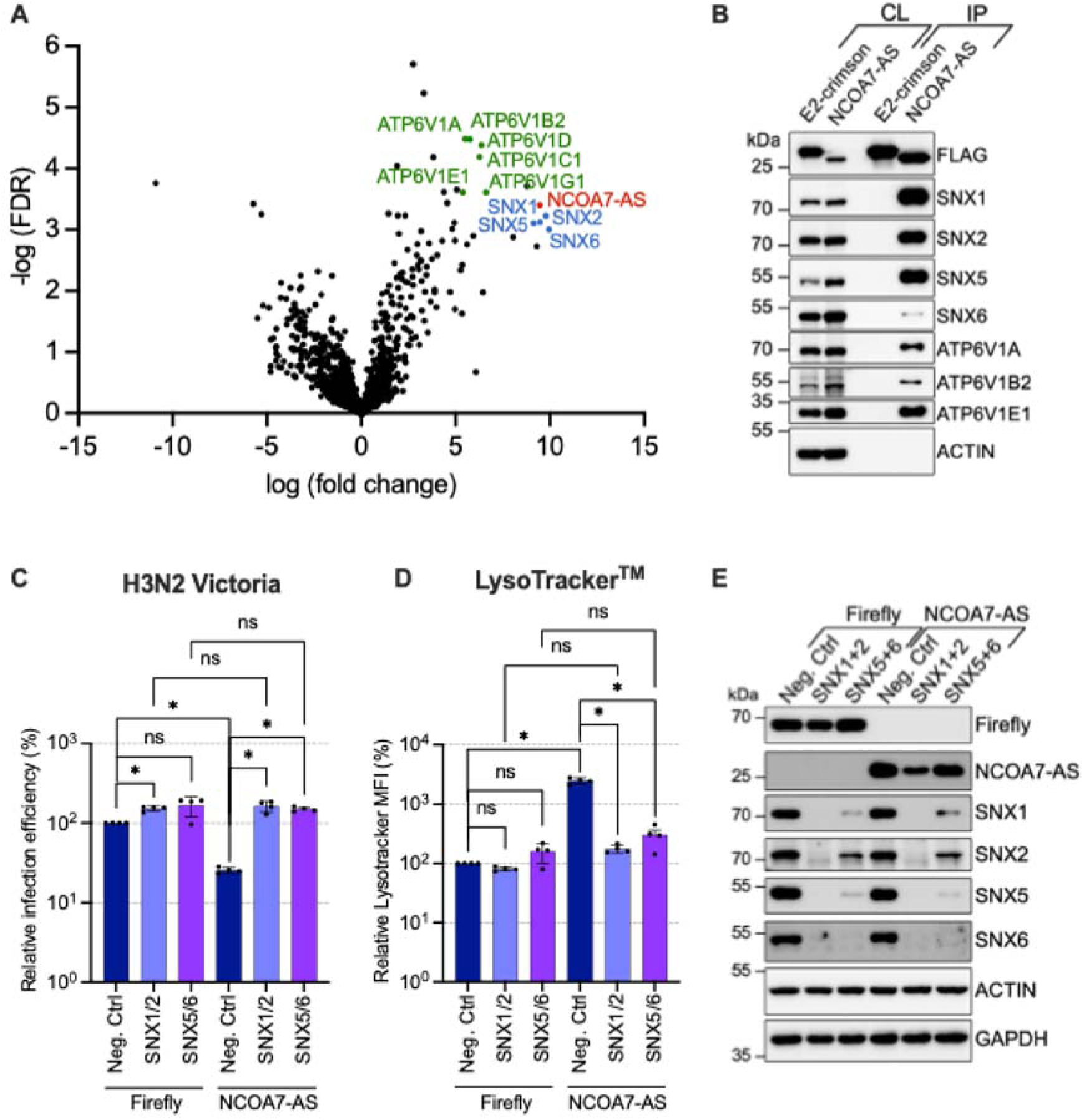
SNX1, SNX2, SNX5, and SNX6 act as physical and functional partners of NCOA7-AS. **A.** Volcano Plot of differentially abundant proteins between NCOA7-AS and control immunoprecipitation, presenting the fold change (in log2) and significance (-Log 10 p value) obtained from a t-test based on intensity values from three independent experiments. **B.** U87-MG cells were transduced with lentiviral vectors expressing FLAG-tagged-NCOA7-AS or -E2-Crimson (a negative control). Cells were lysed and the FLAG-tagged proteins were immunoprecipitated and subsequently analyzed by immunoblotting using the indicated antibodies; Actin served as a loading control. **C-E.** A549 cells were transduced with an all-in-one CRISPRi system, expressing dual non-targeting sgRNAs (Neg. Ctrl) or dual sgRNAs targeting either SNX1/2 or SNX5/6, and then further transduce to express either FLAG-tagged-Firefly or -NCOA7-AS. Cells were infected with IAV (A/Victoria/3/75) encoding a Nanoluciferase reporter gene at an MOI of 0.25 and the relative infection efficiency was measured by monitoring Nanoluciferase activity 8h later (**C**). In parallel, cells were incubated with LysoTracker^TM^ Green DND-26 for 1h at 37°C and the MFI was subsequently analyzed by flow cytometry (**D**). SNX depletion in the cells used in **C-D** was assessed by immunoblotting using the indicated antibodies; Actin and GAPDH served as loading control (**E**). Representative immunoblots (out of 3 independent experiments) are shown (**B, E**) and the mean and SD of three independent experiments (**C-D**); Mann-Whitney test on raw data (**C**) or on log-transformed raw data (**D**).

SNX proteins are important for retrograde transport of proteins between endosomes and the trans-Golgi network (Burd and Cullen, 2014). They notably contain a Phox homology (PX) domain, and some of them are known to interact with phosphoinositides. Hence PX-SNX1 and PX-SNX2 interact with phosphatidylinositol-3-phosphate at the endosomal membrane (Cheever et al., 2001; Yu and Lemmon, 2001; Burda et al., 2002; Gillooly et al., 2003) but not PX-SNX5 and PX-SNX6 (Chandra et al., 2019). Moreover, SNX1/2/5/6 belong to the SNX-BAR subfamily, which possesses a Bin-Amphiphysin-Rvs (BAR) domain. The BAR domain is important for dimerization and to detect and/or induce membrane curvature (Carlton et al., 2004; Frost et al., 2009; Weering et al., 2010). A heterodimer of SNX-BAR proteins, consisting of SNX1 or SNX2 associated with either SNX5 or SNX6, forms the Retromer complex in association with a vacuolar protein sorting (VPS) trimer. The Retromer complex mediates cargo trafficking from the endosome to the trans-Golgi network (Wassmer et al., 2006). However, recent studies have shown that some cargoes could be transported by a SNX1/2-SNX5/6-dependent pathway that was Retromer-independent (Kvainickas et al., 2017; Simonetti et al., 2017; Yong et al., 2020). Interestingly, an additional, non-canonical role has also been reported for SNX5. Indeed, SNX5 has been shown to inhibit several viruses, including IAV, by inducing autophagy at virus-containing endosomes independently of SNX1, SNX2 or the Retromer complex (Dong et al., 2021).

Interactions between NCOA7-AS and SNX1, SNX2, SNX5 and SNX6, identified by mass spectrometry, were validated by co-immunoprecipitation, with all four endogenous SNX proteins co-immunoprecipitating with FLAG-tagged NCOA7-AS-WT but not with the E2-Crimson control in U87-MG cells (Figure 2B).

To assess the role of SNX1, SNX2, SNX5 and SNX6 in NCOA7-AS antiviral activity against IAV and in the overacidification of the endolysosomal compartment, these proteins were depleted in A549 cells using dual CRISPRi targeting SNX1/2 or SNX5/6, alongside a dual non-targeting control (Neg. Ctrl). As mentioned above, SNX1/2 and SNX5/6 function redundantly (Simonetti et al., 2017) and double knockdowns were therefore used to circumvent the redundancy. The depleted cells were transduced with lentiviral vectors to stably express a control (Firefly) or NCOA7-AS. Depletion of either SNX1/SNX2 or SNX5/SNX6 fully restored viral infection in the presence of NCOA7-AS, while they modestly enhanced IAV infection in its absence (Figure 2C). Consistently, the ability of NCOA7-AS to enhance endolysosomal acidification was abolished upon depletion of either SNX1/2 or SNX5/6 (Figure 2D).

Immunoblot analyses confirmed efficient depletion of the targeted proteins (Figure 2E). Notably, each double knockdown affected the stability of the corresponding heterodimer, consistent with previous findings (Simonetti et al., 2017). This effect was more pronounced upon SNX1/2 depletion: while residual SNX1 and SNX2 remained detectable in SNX5/6-depleted cells, SNX5 and SNX6 were no longer detectable in SNX1/2-depleted cells. Moreover, SNX1/2 depletion also impacted the expression levels of Firefly and NCOA7-AS, an effect that was not observed, or only marginal, upon SNX5/6 depletion.

Taken together, these results indicate that these SNX-BAR proteins are required for NCOA7-AS-mediated endolysosomal overacidification and antiviral activity against IAV. However, the contribution of SNX1/2 should be interpreted with caution, as their depletion seemed to affect NCOA7-AS expression levels, whereas the requirement for SNX5/6 appeared more direct.

### The N-terminal domain of NCOA7-AS directly interacts with the Phox domain of SNX5 and SNX6

SNX5 and SNX6 interact with more than 60 cargoes, including cation-independent mannose-6-phosphate receptor (CI-MPR) (Simonetti et al., 2017, 2019) or *Chlamydia trachomatis* IncE (Mirrashidi et al., 2015; Elwell et al., 2017; Paul et al., 2017; Sun et al., 2017). These interactions are mediated by the PX domain of SNX5 or SNX6 (PX-SNX5 and PX-SNX6). Analysis of the sequence requirement to bind SNX5 or SNX6 revealed that cargoes have a conserved secondary structure, consisting of two antiparallel β-strands with one β-strand containing a ΦxΩxΦ motif, where Φ is a hydrophobic residue and Ω, an aromatic residue (Simonetti et al., 2019).

NCOA7-AS contains two main structural domains: the N-terminal domain (NTD) of 53 amino acids and the TLDc domain, the latter being involved in the interaction with the V-ATPase (Eaton et al., 2021). The structure of the human NCOA7-AS TLDc domain has been recently determined (Arnaud-Arnould et al., 2021), but that of NCOA7-AS-NTD was unknown. AlphaFold prediction suggested however that NCOA7-AS-NTD could be organized as two antiparallel β-strands and a small α-helix (Supplementary Figure 1A). Furthermore, the first predicted β-strand contains a ΦxΩxΦ motif (Supplementary Figure 1B), suggesting that NCOA7-AS could directly interact with SNX5 or SNX6.

To test this hypothesis and to map the regions of NCOA7-AS potentially important for SNX binding, the proteins and domains of interest were individually produced in *E. coli* as fusions with Glutathione S-Transferase (GST) or Maltose Binding Protein (MBP) and purified, in order to perform *in vitro* GST pull-down assays. GST-tagged-NCOA7-AS (NCOA7-AS-WT-GST), -NCOA7-AS N-terminal domain (NTD, amino acids 1-53; NCOA7-AS-NTD-GST), -NCOA7-AS TLDc domain (amino acids 54-219, NCOA7-AS-TLDc-GST) and MBP-tagged-PX-SNX1, -PX-SNX5, -PX-SNX6 were used. The GST pull-downs showed that NCOA7-AS interacted with the PX domains of SNX5 and SNX6 but not with the PX domain of SNX1 (Figure 3A). As previously described by other groups, the PX domains of SNX5 and SNX6 contain a unique 38 amino acid insertion, which folds in a helix-turn-helix extension, that is absent in the PX domains of SNX1 and SNX2 (Simonetti et al., 2019). This insertion has already been shown to be important for the interaction with several cargoes, such as CI-MPR (Simonetti et al., 2019) and IncE (Mirrashidi et al., 2015; Elwell et al., 2017; Paul et al., 2017; Sun et al., 2017).

**Figure 3.**
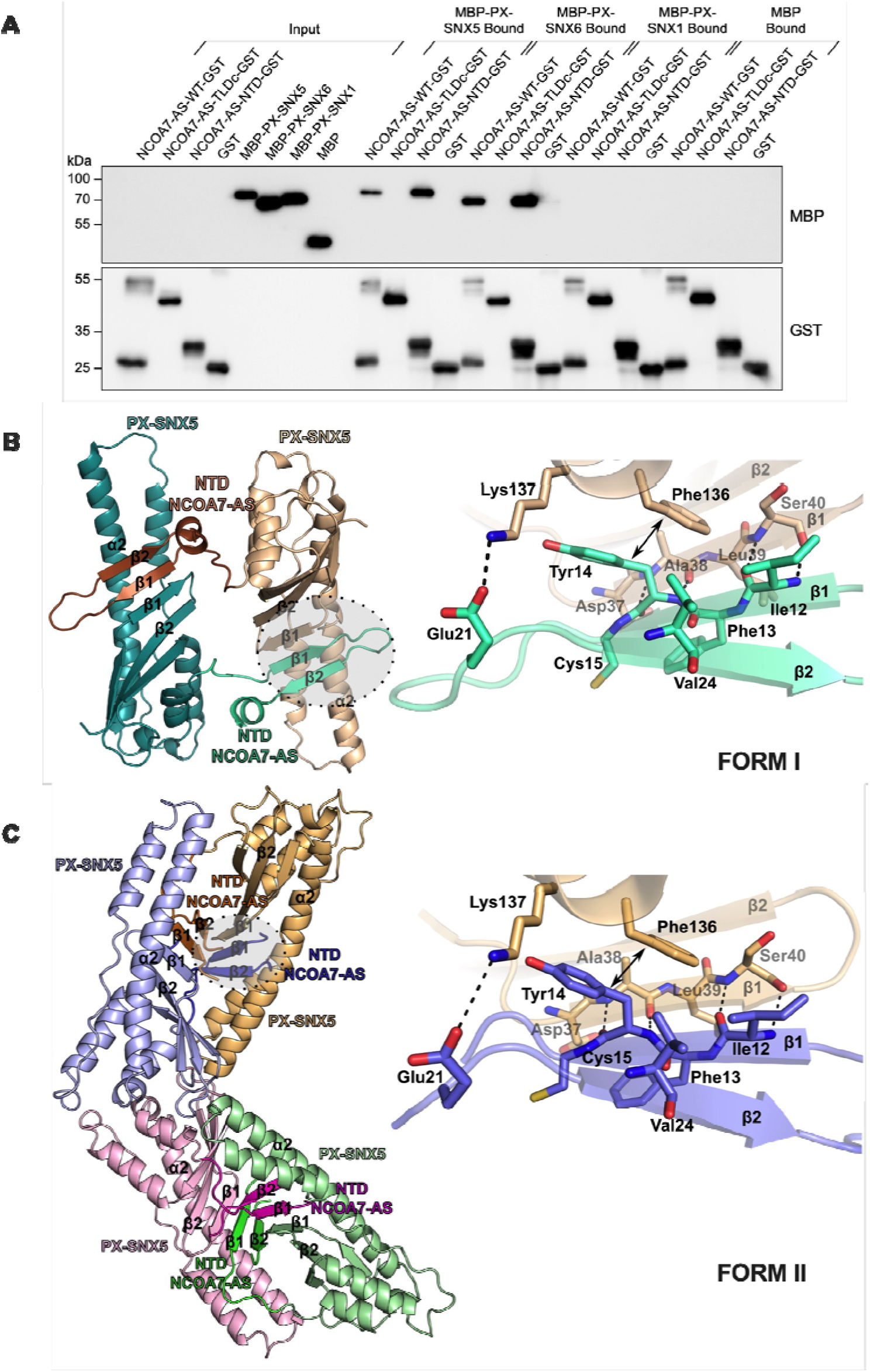
The N-terminal domain of NCOA7-AS interacts directly with the PX domain of SNX5 and SNX6. **A.** Recombinant MBP-tagged-PX-SNX5, -PX-SNX6, -PX-SNX1 or MBP alone, and GST-tagged-NCOA7-AS-WT, -NCOA7-AS-NTD, -NCOA7-AS-TLDc or GST alone, were produced in *Escherichia coli* and purified. GST pull-down assays were performed using these recombinant proteins and analyzed by western blot. **B-C.** Crystal structure of PX-SNX5-NCOA7-AS-NTD chimera. Two crystal forms were solved. In the form I (**B**), the asymmetric unit contains one monomer, the dimer interaction described in here is therefore formed with the symmetric molecule. The second crystal form (**C**) contains four molecules per asymmetric unit and therefore contains two dimers. The overall structures are shown in cartoon representations in **B** and **C**, for all the structures, the NCOA7-AS-NTD is a darker color as compared to the PX-SNX5 domain fused to it. The right panel shows a magnified view of the region, rotated 180° relative to the highlighted zone in grey. The H-bonds interaction between the strands β1 of NCOA7-AS and β1 of the PX domain are shown in black dashed lines. The salt-bridge between Glu21 and Lys137 is also represented by a dashed line. The main interaction between the side chain of Tyr14 from NCOA7-AS and Phe136 of the PX domain is illustrated by the double headed arrows. Val24 also contributes to interact with Phe136 but the interaction is not shown.

Furthermore, the ability of NCOA7-AS to bind to the PX domain of SNX5 and SNX6 was mediated by the NTD and not by the TLDc, as shown by the fact that, contrary to NCOA7-AS-NTD-GST, NCOA7-AS-TLDc-GST was not able to bind PX-SNX5 and PX-SNX6 (Figure 3A). Altogether, these results showed that NCOA7-AS directly interacted with SNX5 and SNX6 PX domains via its N-terminal domain.

In order to better understand the molecular basis of this interaction, we employed a strategy previously described by another team to determine the structure of the two interacting partners (Paul et al., 2017). This strategy consists in designing a chimeric fusion protein with the two binding partners. Such a strategy was successfully used to obtain the crystal structure of PX-SNX5 in complex with a peptide from CI-MPR (Simonetti et al., 2019). Therefore, a chimeric protein composed of PX-SNX5 (amino acids 25-172) and NCOA7-AS N-terminal sequence (amino acids 4-36) was produced. After recombinant expression in *E. coli* and purification (Supplementary Figure 2A), the chimeric protein was crystallized into two different forms (named forms I and II). The X-ray structures of the two distinct crystals forms were solved at high resolution (Figure 3B-C and Supplementary table 2).

PX-SNX5 domain is typically composed of three β-strands (β1, β2 and β3) and three α-helices (α1, α2 and α3) as seen in a previously reported structure (<u>PDB: 5wy2;</u> (Sun et al., 2017)). The N-terminus of NCOA7-AS forms a β-hairpin structure consisting of β1 (residues Gln11-Cys15) and β2 strands (residues Phe23-Ile27) and a short α-helix (residues Val29-Arg35). The β-hairpin structure of NCOA7-AS-NTD interacts with PX-SNX5 by forming an anti-parallel β-sheet. The β1 strand of NCOA7-AS strongly interacts by H-bonds with the β-sheet through β1 of the PX-SNX5 domain, and with its complementary groove at the base of the extended α-helical insertion (α2) of the PX-SNX5 domain. The N-terminal β1 strand of NCOA7-AS forms the primary interface with SNX5, by hydrogen bonds involving the main chain with the β1 strand of the PX-SNX5 domain. The two anti-parallel β-strands of NCOA7-AS are connected by a short loop (Ala16-Pro22) that could not be rebuilt in the crystal form II, however, this loop is not making any direct contact with the SNX5 protein in the other form (form I). The NCOA7-AS β1 strand forms an interface of interaction with the extended α-helical region (α2) of the PX-SNX5 domain. This interaction is mediated by several van der Waals interactions but the strongest interaction is driven by the side chain of Tyr14 from β1 strand of NCOA7-AS, through a hydrophobic interaction with the side chain of Phe136 of SNX5. To a lesser extent, Val24 side chain of NCOA7-AS interacts as well with Phe136 of SNX5. A salt-bridge interaction is also established between PX domain Lys137 and NCOA7-AS Glu21. Of note, although the two crystals forms are packing differently and notably with a different conformation of the NCOA7-AS hairpin in both structures, the β strands interaction and the main interaction between NCOAS-AS Tyr14 and SNX5 Phe136 are conserved in both structures. This is strongly supportive of a specific interaction not driven by the chimera or the crystal packing.

Furthermore, PX-SNX5’s mode of interaction with NCOA7-AS was also compared with the previously solved structures of PX-SNX5 bound to other proteins, namely Sema4C (<u>PDB 6n5z</u>; (Simonetti et al., 2019), CI-MPR (<u>PDB 6n5y</u>; (Simonetti et al., 2019) and IncE (<u>PDB 5wy2</u>; (Sun et al., 2017) (Supplementary Figure 2B). This analysis demonstrated that NCOA7-AS interacted in a similar manner as the other interactors of SNX5, by forming an anti-parallel β-sheet and through a hydrophobic interaction between the Phe136 of PX and either a Tyr or a Phe for the different binding partners. Of note, PX-SNX6 is thought to harbor the same binding specificity to cargoes through its Phe137.

To confirm the importance of these residues for the specific interaction seen in the two crystal structures, Phe136 was mutated into Ala (F136A) on MBP-PX-SNX5. *In vitro* GST pull-down assays with NCOA7-AS-WT-GST demonstrated that PX-SNX5 Phe136 was important for NCOA7-AS binding to SNX5 (Figure 4A). In order to gain more insights into this interaction, the dissociation constant (Kd) between PX-SNX5 domain and the N-terminus of NCOA7-AS was measured by microfluidic diffusional sizing (MDS) (Figure 4B). To do so, the hydrodynamic radius (Rh) of a fluorescently labelled NCOA7-AS peptide corresponding to its NTD was estimated in solution in the absence or the presence of increased concentrations of MBP-PX-SNX5 or MBP-PX-SNX5-F136A domains. In the absence of ligand, NCOA7-AS peptide possesses an Rh of about 1.6 nm, while it reaches 3.1 and 2.7 nm at the highest concentration (250 μM) of WT and mutant PX-SNX5, respectively. The titration curve strongly attests that the single point mutation F136A significantly increases the dissociation constant between PX-SNX5 and NCOA7-AS N-terminal domain (Kd=1.6 µM for the PX-SNX5-WT versus 12.4 µM for the F136A mutant). The analysis of the impact of the reciprocal mutation of Tyr14 to Ala on NCOA7-AS (NCOA7-AS-Y14A) confirmed the importance of this residue for the *in vitro* binding to PX-SNX5 (Figure 4C). Of note, NCOA7-AS Val24 mutation to Ala was also tested. *In vitro* GST pull-down assays showed that the interaction between this mutant and PX-SNX5 was slightly decreased but not abolished (data not shown), contrary to what was observed with the Y14A mutant (Figure 4C).

**Figure 4.**
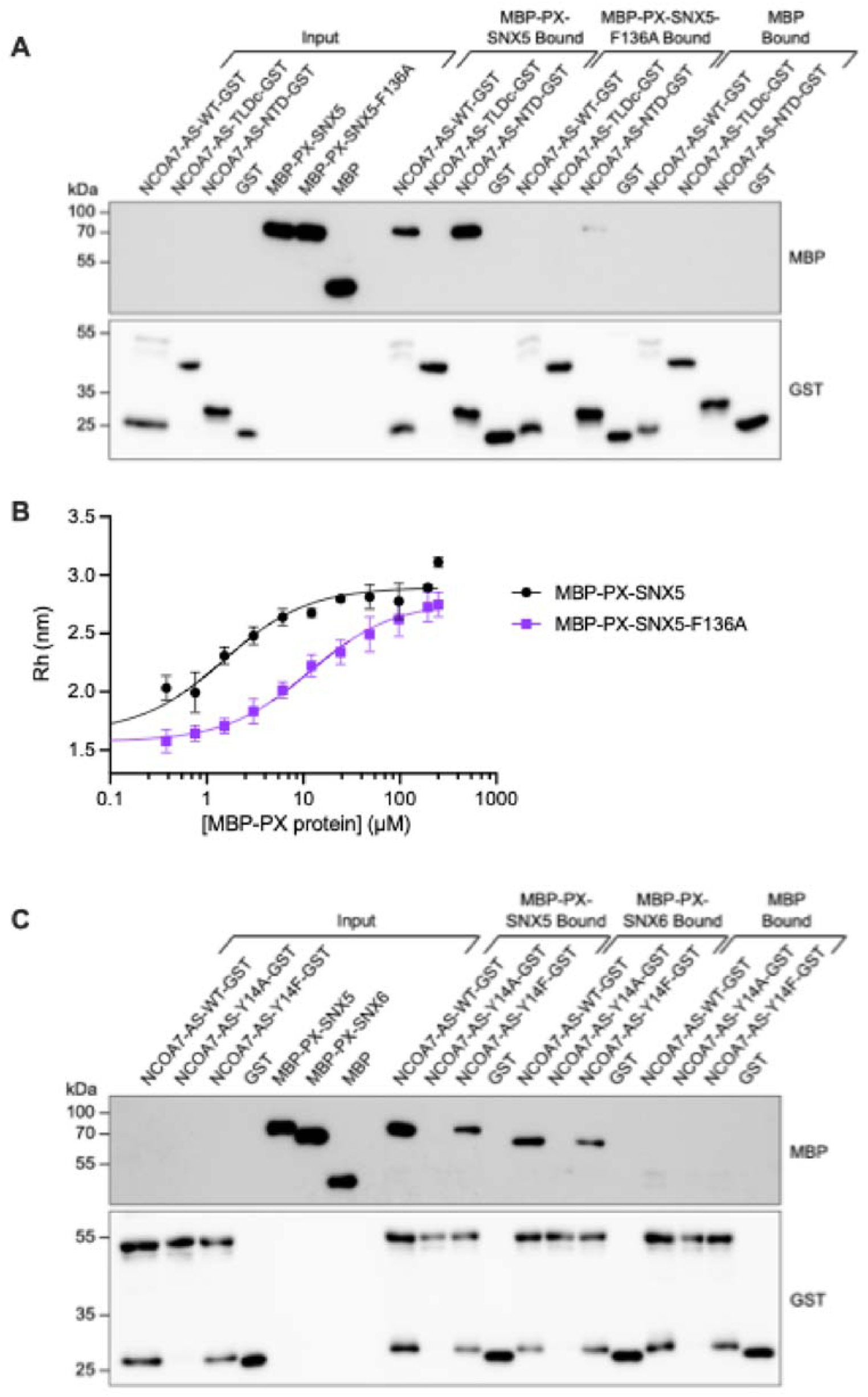
NCOA7-AS-Y14 and SNX5-F136 are important for NCOA7-AS/SNX5 and NCOA7-AS/SNX6 interaction. **A.** Recombinant MBP-tagged-PX-SNX5, -PX-SNX5-F136A or MBP alone, and GST-tagged-NCOA7-AS-WT, -NCOA7-AS-NTD, -NCOA7-AS-TLDc or GST alone were produced in *Escherichia coli* and purified. GST pull-down was performed using these recombinant proteins and analyzed by western blot. **B.** Diffusional sizing of MBP-tagged-PX-SNX5 or -PX-SNX5-F136A with fluorescently labelled NCOA7-AS-NTD (NCOA7-AS-NTD-FITC) was measured using Fluidity One-W. Values represent the means and SD of 3 independent experiments. **C.** Recombinant MBP-tagged-PX-SNX5, -PX-SNX6 or MBP alone and GST-tagged-NCOA7-AS-WT, -NCOA7-AS-Y14A, -NCOA7-AS-Y14F or GST alone were produced in *Escherichia coli* and purified. GST pull-down was performed using these recombinant proteins and analyzed by western blot.

Moreover, Tyr14 was mutated to a minimal conservative Phe substitution, which maintained the aromatic cycle in the side chain. Y14F mutation in NCOA7-AS did not affect its ability to bind to PX-SNX5 and SNX6 (Figure 4C), highlighting the fact that the hydrophobic interaction mediated by Tyr14 and Phe136 drives the interaction between NCOA7-AS and SNX5 or SNX6.

### Residue Tyr14 on NCOA7-AS is essential for NCOA7-AS-mediated acidification of endolysosomes and antiviral activity against IAV

To assess the importance of NCOA7-AS-Y14 in the antiviral activity against IAV, point mutations of NCOA7-AS-Y14 were generated, and A549 cells stably expressing a negative control (Firefly), NCOA7-AS-WT, NCOA7-AS-G91A, NCOA7-AS-Y14A or NCOA7-AS-Y14F were generated. Co-immunoprecipitation experiments first confirmed that NCOA7-AS-Y14A mutant did not interact with any of the 4 SNXs of interest. Minimal conservative NCOA7-AS-Y14F mutant conserved the interaction with SNX5 (and the other SNXs), confirming that the interaction between NCOA7-AS and SNX5 requires π-stacking between NCOA7-AS-Y14 and SNX5-F136 (Figure 5A).

**Figure 5.**
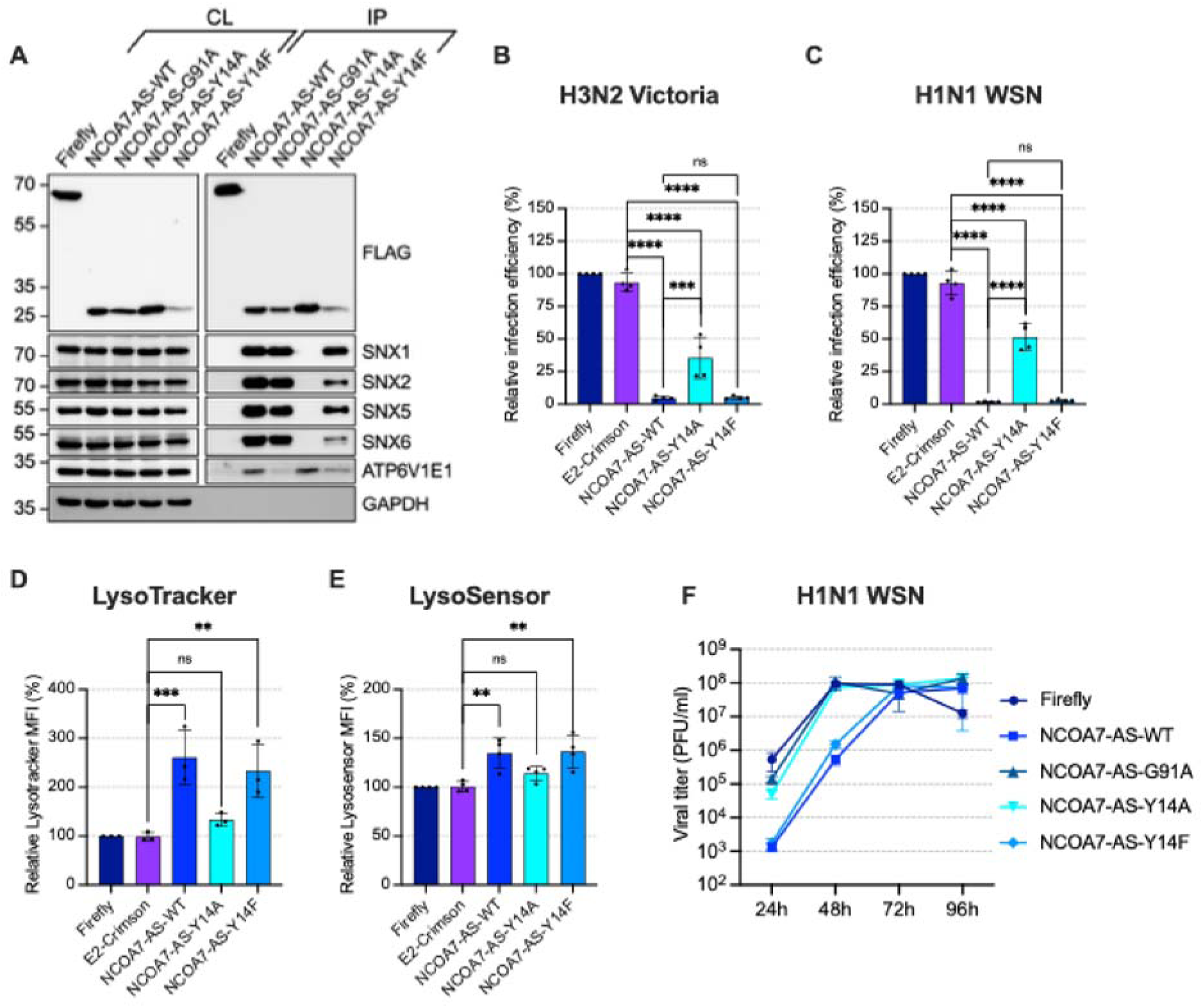
NCOA7-AS Y14 is required for SNX5 binding, antiviral activity against IAV, and endolysosomal overacidification. **A-F.** A549 cells stably expressing FLAG-tagged-Firefly (a control), - FLAG-tagged-E2-crimson (another control), NCOA7-AS-WT, -NCOA7-AS-G91A, -NCOA7-AS-Y14A or -NCOA7-AS-Y14F were generated. **A.** The cells expressing the indicated proteins were lysed and the FLAG-tagged proteins immunoprecipitated, followed by immunoblotting analysis using the indicated antibodies; GAPDH served as a loading control. **B-C.** The cells expressing the indicated proteins were infected with Nanoluciferase-expressing A/Victoria/3/75 at MOI 0.5 (**B**) or A/WSN/1933 at MOI 0.25 (**C**). The relative infection efficiency was measured by monitoring Nanoluciferase activity at 8h post-infection. **D-E.** The cells expressing the indicated proteins were incubated with LysoTracker^TM^ Green DND-26 (**D**) or with LysoSensor^TM^ Green DND-189 (**E**) for 1h at 37°C and the MFI subsequently analyzed by flow cytometry. **F.** The cells expressing the indicated proteins were infected with A/Victoria/3/75 at MOI 0.005 in multicycle replication conditions; the supernatants were harvested at the indicated time points post infection and infectious virus production was measured by plaque assays on MDCK cells. Data represent the mean and SD of 4 (**B-C**) or 3 (**D-E**) independent experiments (one-way ANOVA) or show one representative experiment with the mean and SDs of technical triplicates (**F**).

Although NCOA7-AS-G91A failed to interact with the V-ATPase—as evidenced by the loss of binding to the ATP6V1E1 subunit (Figures 5A and 1B)—it retained interaction with SNX5 and other SNX proteins. This indicates that NCOA7-AS association with SNXs is independent of its interaction with the V-ATPase, consistent with the direct binding observed *in vitro* (Figure 4).

Next, we tested the antiviral activity of these mutants against IAV in single-cycle infection experiments. Compared to NCOA7-AS-WT, NCOA7-AS-Y14A was poorly functional with a significant reduction of antiviral activity against both A/Victoria/3/75 (Figure 5B) and A/WSN/1933 (Figure 5C). In contrast, NCOA7-AS-Y14F retained full antiviral activity despite being less expressed at steady state than NCOA7-AS-WT. The use of LysoTracker^TM^ and LysoSensor^TM^ showed that NCOA7-AS-Y14A had no impact on endolysosomal acidification, whereas NCOA7-AS-Y14F was as active as NCOA7-AS-WT (Figure 5D-E).

To further validate these results, multicycle infection experiments were performed with A/WSN/1933 in A549 cells stably expressing a negative control (Firefly), NCOA7-AS-WT, NCOA7-AS-G91A, NCOA7-AS-Y14A or NCOA7-AS-Y14F (Figure 5F). Both NCOA7-AS-WT and NCOA7-AS-Y14F inhibited IAV infection by 2 logs at 24h and 48h, while NCOA7-AS-G91A and NCOA7-AS-Y14A had no impact on viral growth (similarly to the Firefly negative control). From 72h onwards, the antiviral effect of these proteins was not observed anymore, suggesting that their impact might be transient, saturable, or overcome over time.

Taken together, these data showed that residue Tyr14 on NCOA7-AS, which mediated the interaction with SNX5 and SNX6, was essential for antiviral activity against IAV and for inducing the endolysosomal overacidification.

## Discussion

In this study, we identify key determinants of NCOA7-AS activity against IAV and its impact on endolysosomal pH. Analysis of the NCOA7-AS interactome revealed SNX1, SNX2, SNX5 and SNX6 as novel binding partners, which contributed to NCOA7-AS–mediated overacidification of endolysosomes and antiviral activity. However, the respective contributions of SNX1/2 and SNX5/6 were not equivalent. While depletion of either dimer impaired NCOA7-AS function, interpretation of SNX1/2 involvement was limited by indirect effects on NCOA7-AS expression levels. In contrast, SNX5/6 appeared to play a clearer and more direct role, consistent with their direct interaction with NCOA7-AS.

We were able to characterize in depth, biochemically and structurally, the interaction between SNX5/6 and NCOA7-AS. First, both full length NCOA7-AS and its N-terminal domain interacted directly with the PX domain of SNX5 and SNX6, as shown using recombinant proteins. Such a direct interaction was not observed with SNX1, highlighting the primary role of SNX5/6 in NCOA7-AS function. We solved the X-ray structure of the PX-SNX5 domain bound to the N-terminal domain of NCOA7-AS, which enabled us to map the interface of interaction between the two partners. This showed a binding mode that was similar to the other cargoes of SNX5 (Sun et al., 2017; Simonetti et al., 2019). The N-terminal domain of NCOA7-AS interacted with the PX domain of SNX5 via the formation of a β-sheet with strands β1, β2, and β3 of SNX5 in a complementary groove at the base of the extended α-helical insertion (α2) of the PX-SNX5 domain. This region corresponds to the 38 amino acid insertion in the PX domain that is uniquely present in SNX5 and SNX6, as well as their homologous protein SNX32, but not found in SNX1 and SNX2. This observation presumably explains why we could not detect a direct interaction between NCOA7-AS and SNX1. Notably, the PX domain of SNX32 exhibits a high sequence homology with the PX domains of SNX5 and SNX6. However, SNX32 was not found in the interactome of NCOA7-AS, suggesting that it does not play a role in NCOA7-AS activity. This could be due to subtle differences in SNX32 sequence or subcellular localization as compared to SNX5 and 6, and would require further exploration.

The structure showed that the hydrophobic interaction mediated by SNX5 F136 and NCOA7-AS Y14, facing each other, seemed to be crucial. The essential role of these residues in the NCOA7-AS/SNX5 interaction was validated *in vitro*. The importance of NCOA7-AS Y14 was further confirmed in cells for the binding to SNX5, the overacidification phenotype induced by NCOA7-AS as well as its antiviral activity.

We also identified G91 within the TLDc domain of NCOA7-AS as a critical residue mediating its interaction with the V1 subunits of the V-ATPase. This interaction is essential for both the overacidification of the endosomal compartment and the antiviral activity against IAV. Hence, our current model proposed that ectopic expression of NCOA7-AS disrupts the finely tuned pH gradient of the endolysosomal pathway through its interaction with the V-ATPase, thereby impairing IAV entry. V-ATPase activity is regulated through multiple mechanisms, including the dynamic assembly and disassembly of the V0/V1 complex (reviewed in (Maxson and Grinstein, 2014)). Intriguingly, the Cryo-EM structure of the V-ATPase in complex with another TLDc protein, yeast OXR1, revealed that OXR1 inhibits V-ATPase activity by promoting complex disassembly (Khan et al., 2022). Consistently, the TLDc domains of human OXR1 and NCOA7-AS have also been shown to inhibit V-ATPase activity *in vitro* by inducing disassembly (Oot and Wilkens, 2024). These findings contrast with our own data. However, the *in vitro* study relied on purified, recombinant TLDc domains, which may not fully recapitulate the regulatory complexity present in a cellular context with the full-length proteins. Supporting this, a previous study showed that the long isoform of NCOA7 (NCOA7-FL) enhanced V-ATPase assembly and activity *in vivo* in mice (Castroflorio et al., 2021). Moreover, in human cells, NCOA7-AS ectopic expression clearly increases V-ATPase activity (Doyle et al., 2018) and this study). The apparent discrepancy between *in vitro* and *in vivo*/*ex vivo* data likely reflects the contribution of additional domains in NCOA7-AS and NCOA7-FL that influence the outcome of the TLDc–V-ATPase interaction. In support of this, we found that NCOA7-AS binding partners SNX1/2/5/6 are required for the overacidification of the endolysosomal compartment driven by NCOA7-AS. These findings underscore the importance of binding partners in modulating the activity of TLDc-containing proteins and add an additional layer of complexity to the regulation of V-ATPase function in cells.

Together, these findings shed new light on the molecular mechanism of action of NCOA7-AS, defined novel essential cellular partners, as well as crucial residues for its antiviral activity.

## Acknowlegdements

The authors acknowledge the Paul Scherrer Institute, Villigen, Switzerland for provision of synchrotron radiation beamtime at beamline PXIII-X06DA of the SLS. This work was funded by the National Research Agency (ANR-20-CE15-0010-02-CAIPIRINAS to CG, MB and MW), the European Research Council (ERC) under the European Union’s Horizon 2020 and Horizon Europe’s research and innovation programme (grant agreements 759226, ANTIViR, and `101088622`, InVIRium, to CG), the Institut National de la Santé et de la Recherche Médicale (INSERM) (to CG), a 3-year PhD studentship from the Ministry of Higher Education and Research (to AR), institutional funds from Centre National de la Recherche Scientifique (CNRS) and Montpellier University. OM and CG acknowledge support from the French research network on influenza viruses (ResaFlu; GDR 2073) financed by the CNRS. Finally, the authors acknowledge the flow cytometry and imaging facility MRI, a member of the national infrastructure France-BioImaging supported by the French National Research Agency (previously ANR-10-INBS-04, and now ANR-24-INBS-0005 FBI BIOGEN).

## Author contributions

M.A.A, A.R., M.B. and C.G. designed the study, analyzed the data and wrote the manuscript. M.A.A. carried out the *in vitro* biochemical experiments, molecular cloning and western blots, obtained the crystals as well as analyzed the immunoprecipitation experiments. A.R. did molecular cloning and performed the virology and cellular experiments. M.T. provided technical assistance and together with CG, she prepared the mass spectrometry (MS) samples. S.U. and K.E.K performed the MS analyses. E.R. and M.W. provided feedback and essential reagents. O.M. provided feedback and produced IAV stocks. M.B. supervised the biochemical work and solved the crystal structures. All authors have read and approved the manuscript.

## Materials and methods

### Plasmids

The pRRL.sin.cPPT.SFFV/FLAG-E2-crimson-IRES-PuromycinR.WPRE, pRRL.sin.cPPT.SFFV/FLAG-Firefly-IRES-PuromycinR.WPRE and pRRL.sin.cPPT.SFFV/FLAG-NCOA7-AS-IRES-PuromycinR.WPRE have been previously described (Doyle et al., 2018; Bonaventure et al., 2022). pRRL.sin.cPPT.SFFV/FLAG-NCOA7-AS-G91A-IRES-PuromycinR.WPRE, pRRL.sin.cPPT.SFFV/FLAG-NCOA7-AS-Y14A-IRES-PuromycinR.WPRE and pRRL.sin.cPPT.SFFV/FLAG-NCOA7-AS-Y14F-IRES-PuromycinR.WPRE were obtained by site-directed mutagenesis (overlap-extension PCR).

To express the C-terminal GST-tag version of NCOA7-AS recombinant proteins, a new pET30-MCS-cleavage site for Tobacco Etch Virus (TEV) protease-GST CDS-His6 CDS was created from the pET-30 Ek/LIC expression vector (Novagen). The NCOA7-AS full length, TLDc domain, N-terminal domain of NCOA7-AS, NCOA7-AS-Y14A and NCOA7-AS-Y14F CDS were PCR amplified from the aforementioned pRRL.sin.cPPT.SFFV/IRES-puro.WPRE lentiviral vector plasmid and cloned into the modified pET-30 expression vector using EcoRI and HindIII cloning sites.

For CRISPRi-mediated knockdown, ZIM3 KRAB domain (Alerasool et al., 2020) was fused to the N-terminus of a nuclease-dead Cas9 (dCas9) from plentidCas9-VP64_blast (Addgene, #61425) and cloned into the pLentiCRISPRv2 backbone (Addgene, #52961), where the original sgRNA scaffold was replaced by a new version (Addgene, #133459), generating the pLentiCRISPRi-v3 vector.

CRISPRi LVs coding for sgRNAs targeting SNX1, SNX2, SNX5, SNX6 and control sgRNAs were obtained by cloning annealed oligonucleotides in BsmBI-digested pLentiCRISPRi-v3, as described (Addgene). sgRNAs targeting SNX1, SNX2, SNX5, SNX6, were designed with CRISPick. The sgRNA coding sequences used were as follow: Ctrl neg g#1 5’- AGCACGTAATGTCCGTGGAT-3’, Ctrl neg g#2 5’-CAATCGGCGACGTTTTAAAT-3’, SNX1 g#1 5′- CGGGTGGAAGAAGATGGCGT -3′, SNX1 g#2 5′- TGGTGGTGGCTGTAGCGCTT-3′, SNX2 g#1 5′-CAGCGGAGGAGGTTCCCTCT-3′, SNX2 g#2 5′-GTCCTCCAGATCCTCAAAGT-3′, SNX5 g#1 5′-ACGAGGAGGCCGCCAACCGC-3′, SNX5 g#2 5′-GCGACACGGACGGGAAGCAA-3′, SNX6 g#1 5′-GCGGAGAACACCCACCATCA-3′, SNX6 g#2 5′- ATGATGGTGGGTGTTCTCCG-3′.

For dual CRISPRi constructs, dual sgRNA with SNX1 g#1 and SNX2 g#1 or SNX5 g#1 and SNX6 g#1 or controls (Ctrl neg g#1 and Ctrl neg g#2) were spaced by two BsmBI sites, annealed and cloned into BsmBI-digested pLentiCRISPRi-v3 vector. Then, a sgRNA scaffold CR3/mouse U6 promoter insert was digested with BsmBI and subsequently ligated into BsmBI-digested pLentiCRISPRi-v3 containing the dual sgRNA.

Human SNX1, SNX5 and SNX6 cDNA were amplified by RT-PCR using the SuperScript III™ (Invitrogen) from mRNAs of A549 cells using the following primers: SNX1-forward 5’-TAAGCGGCCGCATGGCGTCGGGTGGTGGTGG -3’; SNX1-reverse 5’- AATTAATTAAGTCGACTTAGGAGATGGCCTTTGCCTCAGG-3; SNX5-forward 5’- GGGCGGCCGCATGGCCGCGGTTCCCGAGTTGCTGC-3’; SNX5-reverse 5’- AATTAATTAAGTCGACTCAGTTATTCTTGAACAAGCTAAT-3’; SNX6-forward 5’- GTGGATCCATGATGGAAGGCCTGGACGACGGCCCGGAC-3’; SNX6-reverse 5’- ACCTCGAGTCATGTGTCTCCATTTAACACTGCC-3’ and were cloned using NotI/SalI for SNX1 and SNX5, BamHI/XhoI for SNX6 in pRRL.sin.cPPT.SFFV/IRES-puro.WPRE.

The PX domain of SNX1, SNX5, and SNX6 were PCR-amplified from the aforementioned pRRL.sin.cPPT.SFFV/IRES-puro.WPRE lentiviral vector plasmids and cloned into pMAL-p4X expression vectors (New England Biolabs) (previously modified using BamHI and HindIII sites to add one KpnI restriction site), using KpnI and HindIII cloning sites. The pMAL-PX-SNX5-F136A was obtained by site-directed mutagenesis (overlap-extension PCR).

The PX-SNX5-beta strands NCOA7-AS CDS was PCR-amplified using an overlapping strategy to generate the chimeric protein and cloned into pET-30 Ek/LIC expression vectors (Novagen) using KpnI and SalI cloning sites. The insert is in frame with the S- and His-tags and a sequence encoding the TEV protease cleavage site was inserted upstream of the chimera coding region to enable tag removal during the protein purification process. All plasmid constructs were verified by Sanger sequencing.

### Cell lines, lentiviral vector production and transduction

Human cell line U87-MG (ARP-2188) were obtained from the AIDS reagent program. Madin-Darby canine kidney (MDCK), A549 cells, and HEK293T were gifts from Prof. W. Barclay (Imperial College London, UK) and Prof. M. Malim (King’s College London, UK), respectively. These cell lines were maintained in Dulbecco’s Modified Eagle Medium (DMEM) (Gibco) supplemented with 10% fetal bovine serum and 100 μg/mL penicillin and 100 units/ml streptomycin (Thermofisher). Lentiviral (LV) stocks were produced as described previously using polyethylenamine (Doyle et al., 2018), either in HEK293T or in HEK293T LentiX (Clontech). Transduction with LVs was performed by incubating cells for 8 h prior to changing media for fresh media. Puromycin (1□μg/mL (Sigma-Aldrich)) and Blasticidin (10 µg/mL (InvivoGen) selections were performed 24h post-transduction.

### IAV production and infection

Nanoluciferase-coding A/WSN/33 (H1N1) and A/Victoria/3/75 (H3N2) (Diot et al., 2016; Doyle et al., 2018) were produced as previously described (Doyle et al., 2018). Briefly, the eight Pol I / PolII plasmids (0.5 μg each) for WSN rescue (a kind gift from Nadia Naffakh and Sandie Munier, Institut Pasteur Paris) or the eight Pol I plasmids (0.5 μg each) and four rescue plasmids (0.32 μg of PB1, PB2, PA and 0.64 μg NP plasmids) for Victoria virus rescue were co-transfected into HEK293T cells in 6-well plate format using Lipofectamine 3000 (Thermo Fisher Scientific). After 24h, the cells were detached and cocultured with MDCK cells in 25 mL flasks for 8h in DMEM containing 10% serum. The medium was then replaced with serum-free medium containing 1 μg/mL of L-1-p-Tosylamino-2-phenylethyl chloromethylketone (TPCK)-treated trypsin. Supernatants were used for virus amplification on MDCK cells. Viral stocks were titrated by plaque assays on MDCK cells.

For IAV single cycle infection, A549 cells stably expressing the constructs of interest were infected at an MOI of 0.5 for A/Victoria/3/75 (H3N2) and at an MOI of 0.25 for A/WSN/1933 for 1h in serum-free DMEM. The media was then replaced with DMEM containing 10% serum, and 8h post-infection, cells were washed with PBS and lysed in Passive Lysis Buffer (Promega). Levels of infection were measured using the Nano-Glo Luciferase Assay System (Promega).

### Growth curve experiments

A549 stably expressing the constructs of interest were infected in triplicate in 12-well plates for 1h at an MOI of 0.005 with A/WSN/1933. Cells were then washed with PBS and incubated in 1mL of serum-free DMEM containing 1 μg/mL of TPCK-treated trypsin. Samples were collected at 24-, 48-, 72- and 96h post-infection and were titrated on MDCK cells by plaque assays.

### Co-immunoprecipitations

U87-MG cells or A549 cells were transduced to stably express single or triple (3x) FLAG-tagged -NCOA7-AS, or mutants, -Firefly or -E2-Crimson. 25 million cells were lysed in lysis buffer (20 mM Hepes-NaOH pH 7.5; 150 mM NaCl; 1% NP-40 and protease inhibitor cocktail (Roche)). Lysates were clarified by centrifugation at 1,000 g for 10 min at 4°C. One fraction of cell lysates, corresponding to the whole-cell lysate was collected at this stage to serve as controls for protein input (10%) and the rest was incubated with FLAG-magnetic beads (Thermo Fisher Scientific) for 4h at 4°C. The beads were washed 5 times in lysis buffer and the immunoprecipitated proteins eluted using 150□μg/mL FLAG peptide (Sigma-Aldrich) in lysis buffer for□2h at 4°C.

### Mass spectrometry analysis

Sample digestion and peptides analysis were essentially performed as previously described (Sirvent et al., 2012). Briefly, purified proteins were loaded on an SDS-PAGE gel. After short migration, each sample was excised in one band. Proteins were digested using trypsin, and obtained peptides were analyzed online by nano-flow HPLC-nanoelectrospray ionization using a Qexactive HF mass spectrometer (Thermo Fisher Scientific) coupled to a nano- LC system (U3000-RSLC, Thermo Fisher Scientific). Desalting and preconcentration of samples were performed on-line on a Pepmap® precolumn (0.3 × 10 mm; Dionex). A gradient consisting of 0–40% B in A for 120 min (A: 0.1% formic acid, 2% acetonitrile in water, and B: 0.1% formic acid in 80% acetonitrile) at 300 nL/min, was used to elute peptides from the capillary reverse-phase column (0.075 × 500 mm, Pepmap®, Dionex). Data were acquired using Xcalibur software (version 4.1). A cycle of one full-scan mass spectrum (375–1,500 m/z) at a resolution of 60,000 (at 200 m/z), followed by 12 data-dependent MS/MS spectra (at a resolution of 30,000, isolation window 1.2 m/z) was repeated continuously throughout the nanoLC separation.

Raw data analysis was performed using the MaxQuant software (version 1.5.5.1) with standard settings. Used database consist of Human entries from Uniprot (reference proteome UniProt 2019_03) and 250 contaminants (MaxQuant contaminant database). Statistical analysis was performed using Perseus (v1.6.1.1). Intensity values were transformed in log2 scale and protein were filtered for at least three valid values in one of the experimental groups (NCOA7-AS or Control). Missing values were imputed by normal distribution (default parameter, assuming that most missing values are missing not at random and represent low abundance measurements). To highlight the proteins most abundant in NCOA7-AS vs Control, we used the built-in Two-sample test function in Perseus using standard parameters (Student’s T-test, permutation base FDR, FDR of 5% and S0 of 0.1).

### Immunoblot analysis

Cells were washed with PBS and were lysed in sample buffer (10 □mM Tris–HCl pH7.6, 150□mM NaCl, 1% Triton X100, 1□mM EDTA, 0.1% deoxycholate, 2% SDS, 5% Glycerol, 100□mM DTT, 0.02% bromophenol blue). Cell lysates were resolved by SDS-PAGE and analyzed by immunoblotting using primary antibodies specific for FLAG (M2, Sigma-Aldrich; 1/1,000), SNX1 (10304-1-AP, Proteintech; 1/1,000), SNX2 (611308, BD Biosciences; 1/1,000), SNX5 (17918-1-AP, Proteintech; 1/1,000), SNX6 (10114-1-AP, Proteintech; 1/300), ATP6V1A (17115-1-AP, Proteintech; 1/1,000), ATP6V1B2 (15097-1- AP, Proteintech; 1/1,000), ATP6V1E1 (15280-1-AP, Proteintech; 1/1,000), actin (A1978, Sigma-Aldrich; 1/5,000), followed by secondary horseradish peroxidase-conjugated anti-mouse or anti-rabbit immunoglobulin antibodies or primary GAPDH antibody coupled to horseradish peroxidase (HRP) GAPDH-HRP antibody (G9295, Sigma-Aldrich, 1/5,000) and primary FLAG antibody coupled to HRP FLAG-HRP antibody (G8592, Sigma-Aldrich, 1/10,000). Chemiluminescence was then measured using the Clarity ECL Western Blotting Substrate, Bio-Rad and a ChemiDoc™ gel imaging system (Bio-Rad).

### Lysosomal compartment expansion and acidification

Transduced A549 cells were incubated with 0.1 μM LysoSensor^TM^ Green DND-189 (ThermoFisher Scientific) or with 0.1 μM LysoTracker^TM^ Green DND-26 (ThermoFisher Scientific) for 1h at 37°C. Cells were then washed, trypsinized and analyzed immediately by flow cytometry on a BD LSRFortessa.

### Protein expression and purification

#### GST tagged proteins

The recombinant plasmids pET-30 Ek/LIC expressing GST, NCOA7-AS-WT-GST and mutants, NCOA7-AS-TLDc-GST, NCOA7-AS-NTD-GST were transformed in an *Escherichia coli* BL21 (DE3) strain resistant to Phage T1 (New England Biolabs) carrying pRARE2. One colony was used to inoculate an overnight culture of 125 mL Lysogeny broth (LB) medium supplemented with kanamycin (50 μg/mL) and chloramphenicol (30 μg/mL). This culture was diluted in 2.5L of LB medium supplemented with the two antibiotics. The cells were grown at 37°C to an optical density at 600 nm of 0.8, then a 30 min cold shock on ice was performed and protein expression was induced with 1 mM Isopropyl-β-D-thiogalactoside (IPTG) and the culture was grown overnight at 16°C (except for GST for which bacteria were grown 3H at 37°C after adding IPTG). The cells were harvested by centrifugation at 8200 *g* for 20 minutes and resuspended in 30 mL of buffer A (50 mM Tris–HCl pH 8, 400 mM NaCl, 5 mM β-mercaptoethanol, 40 mM imidazole, 10% glycerol and 1 mM benzamidine). The cells were disrupted by sonication and cell debris were removed by centrifugation at 28 000 *g* for 60 minutes. The supernatant was loaded at 4°C on Ni–NTA sepharose beads (GE17-5318-01, Sigma-Aldrich) previously equilibrated with buffer A. The beads were washed once with buffer A and twice with buffer B (50 mM Tris–HCl pH 8, 1 M NaCl, 5 mM β-mercaptoethanol, 40 mM imidazole and 10% glycerol) and elution was performed with buffer E (50 mM Tris–HCl pH 8, 600 mM NaCl, 5 mM β-mercaptoethanol, 250 mM imidazole, 10% glycerol). The eluted protein was dialysed (dialysis-bag cutoff 12–15 kDa) against 1L dialysis buffer D (50 mM Tris–HCl pH 8, 600 mM NaCl, 5 mM β-mercaptoethanol, 10% glycerol) overnight at 4°C. After dialysis, the proteins were concentrated to 5 mg/mL using a Vivaspin® column (10 kDa cutoff, Sartorius), loaded onto a size-exclusion chromatography column (Superdex 200 increase 10/300 GL, GE Healthcare) and eluted with buffer F (50 mM Tris–HCl pH 8, 600 mM NaCl, 5 mM β-mercaptoethanol, 10% glycerol). Aliquots of purified GST-tagged proteins were snap frozen in liquid nitrogen and stored at −80 °C.

#### MBP-tagged proteins

The recombinant plasmids pMAL-p4X expressing MBP-PX-SNX1, 5, and 6 were transformed in an *Escherichia coli* BL21 (DE3) strain resistant to Phage T1 (New England Biolabs) carrying pRARE2. One colony was used to inoculate an overnight culture of 125mL Lysogeny broth (LB) medium supplemented with kanamycin (50 μg/mL) and chloramphenicol (30 μg/mL). This culture was diluted in 2.5L of LB medium supplemented with the two antibiotics. The cells were grown at 37°C to an optical density at 600 nm of 0.8, then a 60 min cold shock on ice was performed, protein expression was induced with 1 mM IPTG and the culture was grown overnight at 16°C. The cells were harvested by centrifugation at 8 200 *g* for 20 minutes and resuspended in 30 mL of buffer A (50 mM Tris–HCl pH 8, 400 mM NaCl, 5 mM β-mercaptoethanol, 10% glycerol and 1 mM benzamidine). The cells were disrupted by sonication and cell debris were removed by centrifugation at 28 000 *g* for 60 minutes. The supernatant was loaded at 4°C on Amylose resin previously equilibrated with buffer A. The beads were washed once with buffer A and twice with buffer B (50 mM Tris–HCl pH 8, 1 M NaCl, 5 mM β-mercaptoethanol and 10% glycerol) and elution was performed with buffer E (50 mM Tris–HCl pH 8, 600 mM NaCl, 5 mM β-mercaptoethanol, 20 mM maltose, 10% glycerol). The eluted protein was dialysed (dialysis-bag cutoff 12–15 kDa) against 1L dialysis buffer D (50 mM Tris–HCl pH 8, 600 mM NaCl, 5 mM β-mercaptoethanol, 10% glycerol) overnight at 4°C. After dialysis, the proteins were concentrated to 5 mg/mL using a Vivaspin® column (10 kDa cutoff, Sartorius), loaded onto a size-exclusion chromatography column (Superdex 200 increase 10/300 GL, GE Healthcare) and eluted with buffer F (50 mM Tris–HCl pH 8, 600 mM NaCl, 5 mM β-mercaptoethanol, 10% glycerol). Aliquots of purified MBP-tagged proteins were snap frozen in liquid nitrogen and stored at −80°C.

### GST pull-down

Recombinant GST-tagged proteins (14 to 35 µg) were incubated 2h at 4°C with glutathione agarose beads prewashed in pull-down assay buffer (50 mM Tris–HCl pH 8, 600 mM NaCl, 5 mM β-mercaptoethanol, 10% glycerol). Beads were then washed 3 times with pull-down assay buffer, then incubated for 2h at 4°C with 85 µg of recombinant MBP-tagged protein preincubated with glutathione agarose beads for 2h at 4°C. Beads were finally washed 3 times with pull-down assay buffer before adding 25 µl of Laemmli 2X. Samples were resolved by SDS-PAGE on a 10% acrylamide gel after 10 min boiling. Immunoblotting was performed by incubation with a primary GST antibody coupled to horseradish peroxidase (HRP) (A7340, Sigma-Aldrich) and an MBP antibody (E8032, NEB) followed by an HRP-conjugated secondary antibody. Bioluminescence was then measured (Clarity ECL Western Blotting Substrate, Bio-Rad) using a ChemiDoc system (Bio-Rad).

### *K*d determination

Recombinant MBP-PX-SNX5 and MBP-PX-SNX5-F136A were dialysed overnight at 4°C against 1L of PBS-10% glycerol. Concentrated proteins (0.38 µM to 250 µM) were mixed with 57 nM final FITC conjugated NCOA7-AS peptide (NH2-RLPLDIQIFY CARPDEEPFV KIITVEEAKR R-COOH) before loading on a Fluidity One chip. Hydrodynamic Radius were given by Fluidity One-W apparatus (Fluidic Analytics) and used to calculate *K*d using the equation: 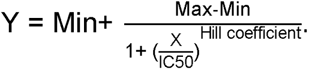.

### Expression and purification of the chimera PX-SNX5-beta strands NCOA7-AS

The recombinant plasmid pET-30-PX-SNX5-beta strands NCOA7-AS was transformed in an *Escherichia coli* BL21 (DE3) strain resistant to Phage T1 (New England Biolabs) carrying pRARE2. One colony was used to inoculate an overnight culture of 125mL Lysogeny broth (LB) medium supplemented with kanamycin (50 μg/mL) and chloramphenicol (30 μg/mL). This culture was diluted in 2.5L of LB medium supplemented with the two antibiotics. The cells were grown at 37°C to an optical density at 600 nm of 0.8, then protein expression was induced with 1 mM IPTG and the culture was grown 3H at 37°C. The cells were harvested by centrifugation at 8 200 *g* for 20 minutes and resuspended in 30 mL of buffer A (50 mM Tris–HCl pH 8, 400 mM NaCl, 5 mM β-mercaptoethanol, 40 mM imidazole, 10% glycerol and 1 mM benzamidine). The cells were disrupted by sonication and cell debris were removed by centrifugation at 28 000 *g* for 60 minutes. The supernatant was loaded at 4°C on Ni–NTA sepharose beads previously equilibrated with buffer A. The beads were washed once with buffer A and twice with buffer B (50 mM Tris–HCl pH 8, 1 M NaCl, 5 mM β-mercaptoethanol, 40 mM imidazole and 10% glycerol) and elution was performed with buffer E (50 mM Tris–HCl pH 8, 200 mM NaCl, 5 mM β-mercaptoethanol, 250 mM imidazole, 10% glycerol). The eluted protein was incubated with His-tagged TEV protease purified in our laboratory in a 1:100 (w:w) ratio; the cleavage reaction was performed during dialysis (dialysis-bag cutoff 12–15 kDa) against 1 L dialysis buffer D (50 mM Tris–HCl pH 8, 200 mM NaCl, 5 mM β-mercaptoethanol) overnight at 4 °C. After dialysis, the proteins were centrifuged for 20 min at 28 000 *g* and the supernatant was loaded again at 4 °C on Ni–NTA sepharose beads equilibrated with buffer D. The proteins without the His-tag were collected in the flow-through, concentrated to 5 mg/ml using a Vivaspin column (10 kDa cutoff, Sartorius), loaded onto a size-exclusion chromatography column (Superdex 75 Increase 10/300 GL, GE Healthcare), and eluted with buffer F (50 mM Tris–HCl pH 8, 100 mM NaCl, 5 mM β-mercaptoethanol). Following this protocol, 3.5 mg pure protein was obtained from 1L culture.

### Protein crystallization

Crystals of the trigonal form were obtained in sitting drops using the SwissCi (MRC) 96 well plate mixing 0.6 μL of protein solution concentrated to 17 mg/mL with 0.6 μL of reservoir solution made of 0.2 M Calcium acetate hydrate, 0.1M Sodium Cacodylate pH6.5 and 18% w/v polyethylene glycol 8000 and equilibrated against a reservoir volume of 50 μL at 18°C. The monoclinic crystals were obtained similarly by mixing 0.6 μL of protein solution concentrated to 17 mg/mL with 0.6 μL of reservoir solution made of 0.1M sodium citrate tribasic dihydrate pH 5.0, 30% v/v jeffamine® ED-2001 pH 7.0 and were not cryoprotected before being cryocooled.

### X-ray Data collection and processing

Data collection was performed on the PXIII-X06DA beamline at the Swiss Light Source, Paul Scherrer Institute, Villigen, Switzerland. Data were processed and scaled with the XDS package (Kabsch, 2010). The phase problem was solved by molecular replacement using Phaser (McCoy, 2007) and the Phenix package (Liebschner et al., 2019), and the structure of the human SNX5 PX domain (5wy2) (Sun et al., 2017) as a search template. The structures were further manually rebuilt with Coot (Casañal et al., 2020) and refined with the Phenix package (Liebschner et al., 2019).

### Statistical analyses

Statistical analyses were performed using GraphPad Prism software using the indicated tests. p-values are indicated as the following: ns = not significant; p < 0.05 = *; p < 0.01 = **; p < 0.001 = ***; and p < 0.0001 = ****.

## Supplementary Figures

**Supplementary Figure 1.**
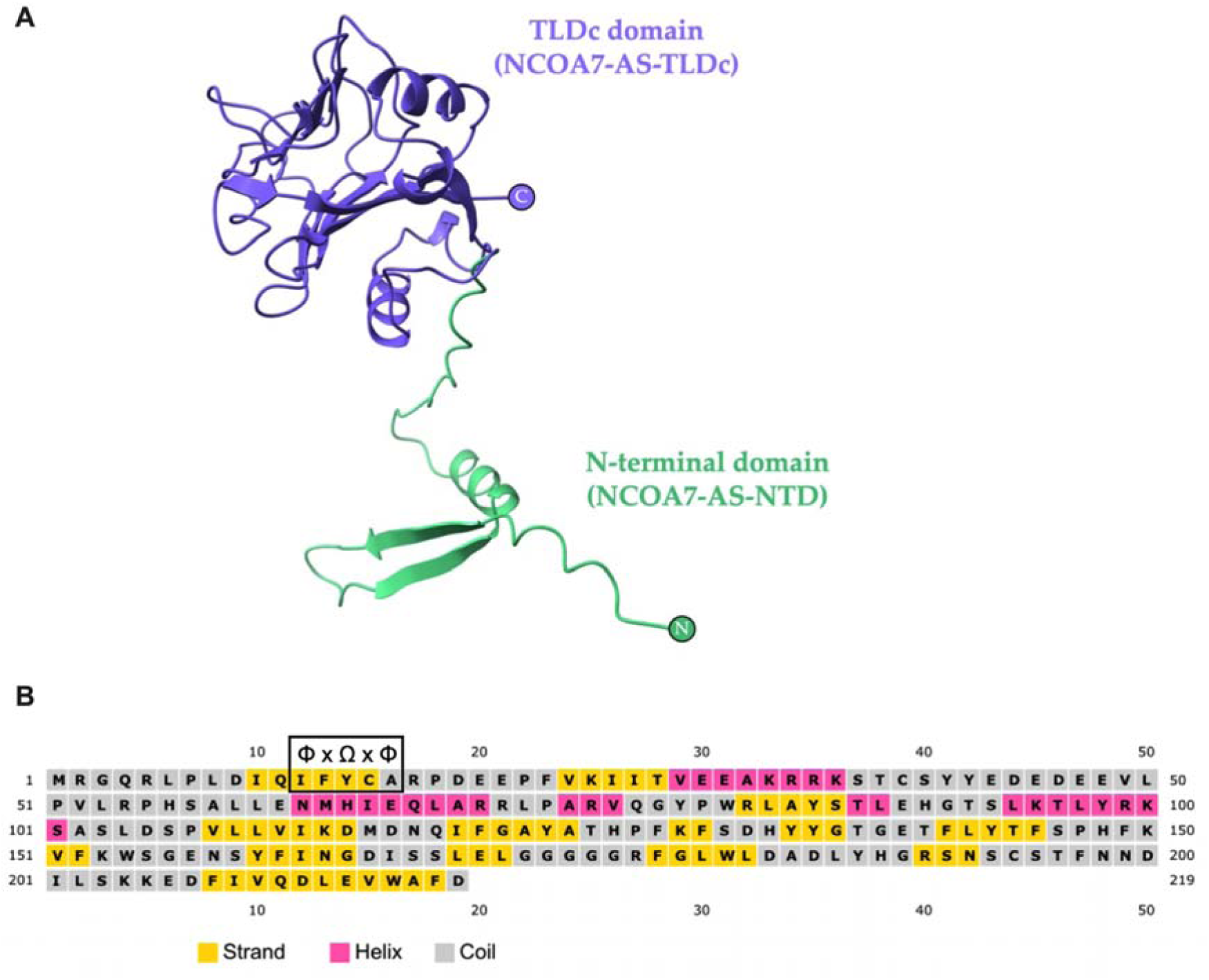
**A.** AlphaFold prediction of NCOA7-AS with the N-terminal domain (NTD-amino-acids 1-53) colored in green and the C-terminal TLDc colored in purple. The NTD contains two antiparallel β-strands and a small α-helix. **B.** PSIPRED predictions, highlighting the presence of a ΦxΩxΦ (black box) in the first β-strand of NCOA7-AS.

**Supplementary Figure 2.**
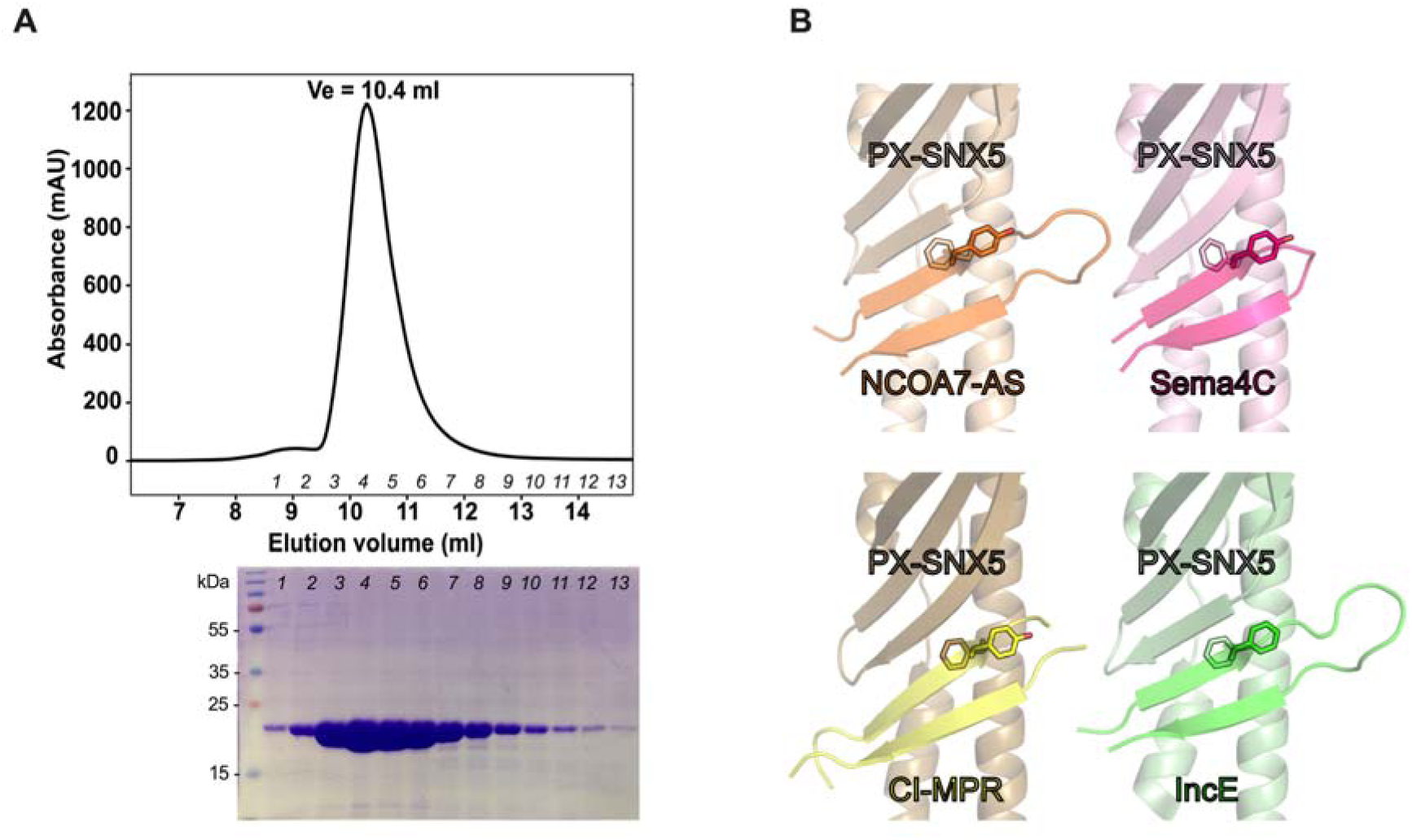
**A.** The recombinant PX-SNX5-NCOA7-AS chimeric protein was produced in *Escherichia coli* and purified. The elution profile on Superdex 75 column is shown with the different fractions collected that have been analyzed by Coomassie staining. **B.** Comparison of the interface of PX-SNX5/NCOA7-AS with all the previously solved structure of PX-SNX5 bound to other binding partners. PX-SNX5::Sema4C (PDB: 6n5z), PX-SNX5::CI-MPR (PDB: 6n5y) and PX-SNX5::IncE (PDB: 5wy2).

## Supplementary Tables

**Supplementary Table 1.**
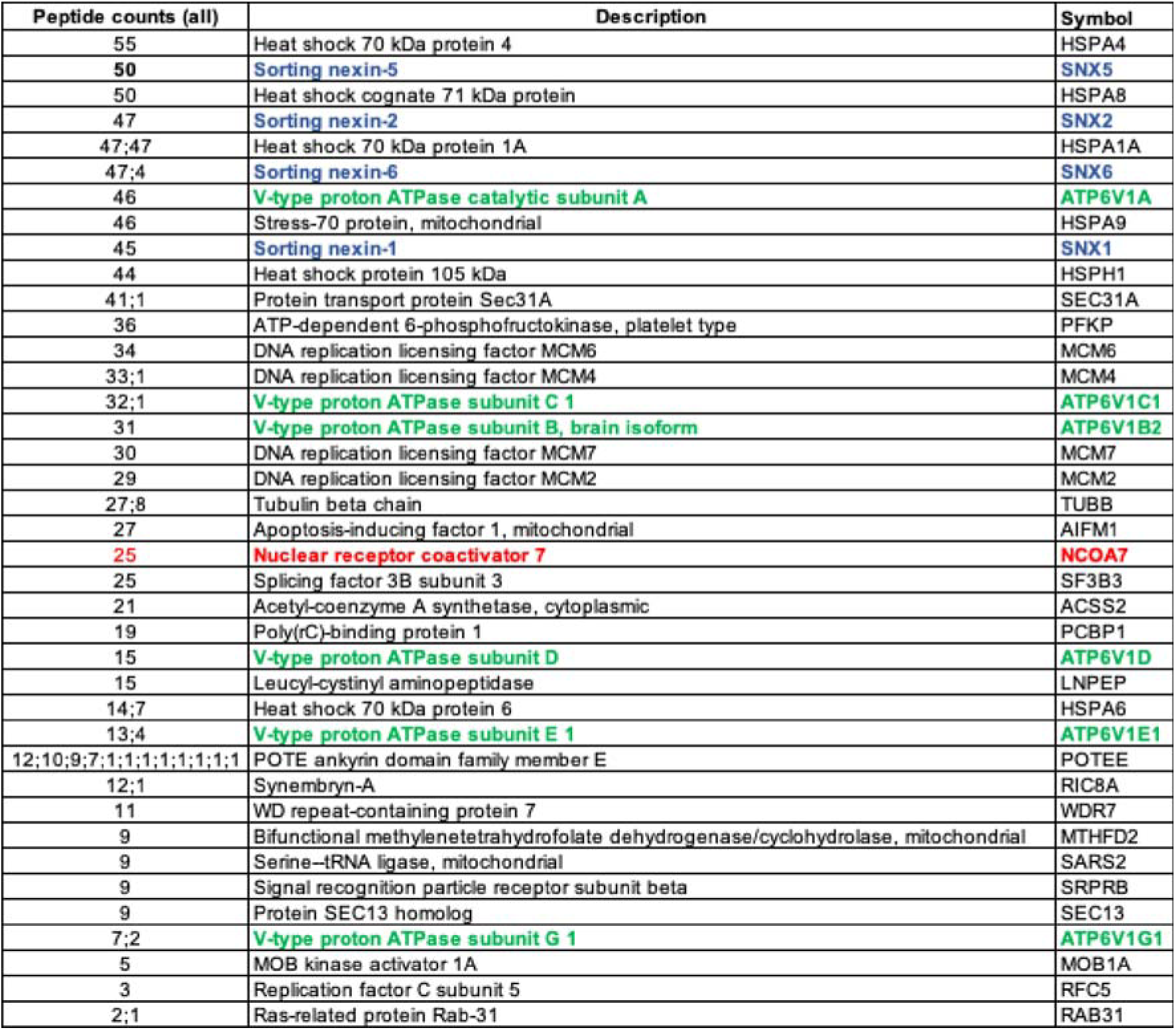
Mass Spectrometry data. Detailed xlsx spreadsheet to download. Summary Table:

| Peptide counts (all) | Description | Symbol |
| --- | --- | --- |
| 55 | Heat shock 70 kDa protein 4 | HSPA4 |
| 50 | Sorting nexin-5 | SNX5 |
| 50 | Heat shock cognate 71 kDa protein | HSPA8 |
| 47 | Sorting nexin-2 | SNX2 |
| 47;47 | Heat shock 70 kDa protein 1A | HSPA1A |
| 47;4 | Sorting nexin-6 | SNX6 |
| 46 | V-type proton ATPase catalytic subunit A | ATP6V1A |
| 46 | Stress-70 protein, mitochondrial | HSPA9 |
| 45 | Sorting nexin-1 | SNX1 |
| 44 | Heat shock protein 105 kDa | HSPH1 |
| 41;1 | Protein transport protein Sec31A | SEC31A |
| 36 | ATP-dependent 6-phosphofructokinase, platelet type | PFKP |
| 34 | DNA replication licensing factor MCM6 | MCM6 |
| 33;1 | DNA replication licensing factor MCM4 | MCM4 |
| 32;1 | V-type proton ATPase subunit C 1 | ATP6V1C1 |
| 31 | V-type proton ATPase subunit B, brain isoform | ATP6V1B2 |
| 30 | DNA replication licensing factor MCM7 | MCM7 |
| 29 | DNA replication licensing factor MCM2 | MCM2 |
| 27;8 | Tubulin beta chain | TUBB |
| 27 | Apoptosis-inducing factor 1, mitochondrial | AIFM1 |
| 25 | Nuclear receptor coactivator 7 | NCOA7 |
| 25 | Splicing factor 3B subunit 3 | SF3B3 |
| 21 | Acetyl-coenzyme A synthetase, cytoplasmic | ACSS2 |
| 19 | Poly(rC)-binding protein 1 | PCBP1 |
| 15 | V-type proton ATPase subunit D | ATP6V1D |
| 15 | Leucyl-cystinyl aminopeptidase | LNPEP |
| 14;7 | Heat shock 70 kDa protein 6 | HSPA6 |
| 13;4 | V-type proton ATPase subunit E 1 | ATP6V1E1 |
| 12;10;9;7;1;1;1;1;1;1;1 | POTE ankyrin domain family member E | POTEE |
| 12;1 | Synembryn-A | RIC8A |
| 11 | WD repeat-containing protein 7 | WDR7 |
| 9 | Bifunctional methylenetetrahydrofolate dehydrogenase/cyclohydrolase, mitochondrial | MTHFD2 |
| 9 | Serine-tRNA ligase, mitochondrial | SARS2 |
| 9 | Signal recognition particle receptor subunit beta | SRPRB |
| 9 | Protein SEC13 homolog | SEC13 |
| 7;2 | V-type proton ATPase subunit G 1 | ATP6V1G1 |
| 5 | MOB kinase activator 1A | MOB1A |
| 3 | Replication factor C subunit 5 | RFC5 |
| 2;1 | Ras-related protein Rab-31 | RAB31 |

**Supplementary Table 2.** X-ray data collection and refinement statistics.

|  | **Nt-NCOA7-PX-SNX5**<br>**Form I** | **Nt-NCOA7-PX-SNX5**<br>**Form II** |
| --- | --- | --- |
| **PDB deposition code** | 9IAE | 9IGY |
| **Wavelength (Å)** | 1 | 1 |
| **Resolution range** | 48.1 - 2.3 (2.38 - 2.3) | 46.1 - 2.3 (2.36 - 2.3) |
| **Space group** | *P* 3<sub>2</sub> 2 1 | *P* 1 2<sub>1</sub> 1 |
| **Unit cell** | 63.13 63.13 101.51<br>90 90 120 | 46.9 76.99 110.85<br>90 100.181 90 |
| **Total reflections** | 210470 (25566) | 230324 (29031) |
| **Multiplicity** | 19.6 (20.14) | 6.8 (7.0) |
| **Completeness (%)** | 98.6 (100.00) | 97.6 (99.9) |
| **Mean I/sigma(I)** | 16.02 (2.23) | 10.98 (1.61) |
| **Wilson B-factor** | 37.3 | 38.6 |
| **R-meas (%)** | 19.3 (166.3) | 16.2 (130.1) |
| **CC1/2** | 0.999 (0.755) | 0.997 (0.695) |
| **Reflections used in refinement** | 10727 (1332) | 33853 (2487) |
| **Reflections used for R-free** | 1076 (106) | 2015 (253) |
| **R-work** | 0.201 (0.247) | 0.227 (0.289) |
| **R-free** | 0.239 (0.346) | 0.262 (0.323) |
| **Number of non-hydrogen atoms** | 1595 | 5665 |
| **macromolecules** | 1434 | 5437 |
| **solvent** | 161 | 228 |
| **Protein residues** | 173 | 665 |
| **RMSD (bonds, Å)** | 0.006 | 0.002 |
| **RMSD (angles, °)** | 0.52 | 0.41 |
| **Ramachandran favored (%)** | 98.21 | 97.53 |
| **Ramachandran allowed (%)** | 1.79 | 2.47 |
| **Ramachandran outliers (%)** | 0.00 | 0.00 |
| **Rotamer outliers (%)** | 1.25 | 0.81 |
| **Clashscore** | 3.16 | 3.24 |
| **Average B-factor (Å<sup>2</sup>)** | 46.13 | 46.58 |
| **macromolecules** | 44.91 | 46.75 |
| **solvent** | 45.37 | 42.60 |

Statistics for the highest-resolution shell are shown in parentheses.

## References

Alerasool, N., Segal, D., Lee, H., Taipale, M., 2020. An efficient KRAB domain for CRISPRi applications in human cells. Nat. Methods 17, 1093–1096. 10.1038/s41592-020-0966-x

Arnaud-Arnould, M., Tauziet, M., Moncorgé, O., Goujon, C., Blaise, M., 2021. Crystal structure of the TLDc domain of human NCOA7-AS. Acta Crystallogr F Struct Biol Commun 77, 230–237. 10.1107/s2053230x21006853

Blaise, M., Alsarraf, H.M.A.B., Wong, J.E.M.M., Midtgaard, S.R., Laroche, F., Schack, L., Spaink, H., Stougaard, J., Thirup, S., 2012. Crystal structure of the TLDc domain of oxidation resistance protein 2 from zebrafish. Proteins 80, 1694–1698. 10.1002/prot.24050

Bonaventure, B., Rebendenne, A., Valadão, A.L.C., Arnaud-Arnould, M., Gracias, S., Gracia, F.G. de, McKellar, J., Labaronne, E., Tauziet, M., Vivet-Boudou, V., Bernard, E., Briant, L., Gros, N., Djilli, W., Courgnaud, V., Parrinello, H., Rialle, S., Blaise, M., Lacroix, L., Lavigne, M., Paillart, J.-C., Ricci, E.P., Schulz, R., Jouvenet, N., Moncorgé, O., Goujon, C., 2022. The DEAD box RNA helicase DDX42 is an intrinsic inhibitor of positive-strand RNA viruses. EMBO Rep 23, e54061. 10.15252/embr.202154061

Burd, C., Cullen, P.J., 2014. Retromer: A Master Conductor of Endosome Sorting. Cold Spring Harbor Perspectives in Biology 6, a016774–a016774. 10.1101/cshperspect.a016774

Burda, P., Padilla, S.M., Sarkar, S., Emr, S.D., 2002. Retromer function in endosome-to-Golgi retrograde transport is regulated by the yeast Vps34 PtdIns 3-kinase. J Cell Sci 115, 3889–3900. 10.1242/jcs.00090

Carlton, J., Bujny, M., Peter, B.J., Oorschot, V.M.J., Rutherford, A., Mellor, H., Klumperman, J., McMahon, H.T., Cullen, P.J., 2004. Sorting nexin-1 mediates tubular endosome-to-TGN transport through coincidence sensing of high- curvature membranes and 3-phosphoinositides. Curr Biol 14, 1791–1800. 10.1016/j.cub.2004.09.077

Casañal, A., Lohkamp, B., Emsley, P., 2020. Current developments in *Coot* for macromolecular model building of Electron Cryo-microscopy and Crystallographic Data. Protein Science 29, 1055–1064. 10.1002/pro.3791

Castroflorio, E., Hoed, J. den, Svistunova, D., Finelli, M.J., Cebrian-Serrano, A., Corrochano, S., Bassett, A.R., Davies, B., Oliver, P.L., 2021. The Ncoa7 locus regulates V-ATPase formation and function, neurodevelopment and behaviour. Cell Mol Life Sci 78, 3503–3524. 10.1007/s00018-020-03721-6

Chandra, M., Chin, Y.K.-Y., Mas, C., Feathers, J.R., Paul, B., Datta, S., Chen, K.-E., Jia, X., Yang, Z., Norwood, S.J., Mohanty, B., Bugarcic, A., Teasdale, R.D., Henne, W.M., Mobli, M., Collins, B.M., 2019. Classification of the human phox homology (PX) domains based on their phosphoinositide binding specificities. Nat. Commun. 10, 1528. 10.1038/s41467-019-09355-y

Cheever, M.L., Sato, T.K., Beer, T. de, Kutateladze, T.G., Emr, S.D., Overduin, M., 2001. Phox domain interaction with PtdIns(3)P targets the Vam7 t-SNARE to vacuole membranes. Nat Cell Biol 3, 613–618. 10.1038/35083000

Cotter, K., Stransky, L., McGuire, C., Forgac, M., 2015. Recent Insights into the Structure, Regulation, and Function of the V-ATPases. Trends in Biochemical Sciences 40, 611–622. 10.1016/j.tibs.2015.08.005

Diot, C., Fournier, G., Santos, M.D., Magnus, J., Komarova, A., Werf, S.V.D., Munier, S., Naffakh, N., 2016. Influenza A Virus Polymerase Recruits the RNA Helicase DDX19 to Promote the Nuclear Export of Viral mRNAs. Sci Rep 6, 33763. 10.1038/srep33763

Dong, X., Yang, Y., Zou, Z., Zhao, Y., Ci, B., Zhong, L., Bhave, M., Wang, L., Kuo, Y.-C., Zang, X., Zhong, R., Aguilera, E.R., Richardson, R.B., Simonetti, B., Schoggins, J.W., Pfeiffer, J.K., Yu, L., Zhang, X., Xie, Y., Schmid, S.L., Xiao, G., Gleeson, P.A., Ktistakis, N.T., Cullen, P.J., Xavier, R.J., Levine, B., 2021. Sorting nexin 5 mediates virus-induced autophagy and immunity. Nature 589, 456–461. 10.1038/s41586-020-03056-z

Dou, D., Revol, R., Östbye, H., Wang, H., Daniels, R., 2018. Influenza A Virus Cell Entry, Replication, Virion Assembly and Movement. Front. Immunol. 9, 1581. 10.3389/fimmu.2018.01581

Doyle, T., Moncorgé, O., Bonaventure, B., Pollpeter, D., Lussignol, M., Tauziet, M., Apolonia, L., Catanese, M.-T., Goujon, C., Malim, M.H., 2018. The interferon-inducible isoform of NCOA7 inhibits endosome-mediated viral entry. Nat Microbiol 3, 1369–1376. 10.1038/s41564-018-0273-9

Eaton, A.F., Brown, D., Merkulova, M., 2021. The evolutionary conserved TLDc domain defines a new class of (H+)V-ATPase interacting proteins. Sci Rep 11, 22654. 10.1038/s41598-021-01809-y

Elwell, C.A., Czudnochowski, N., Dollen, J.V., Johnson, J.R., Nakagawa, R., Mirrashidi, K., Krogan, N.J., Engel, J.N., Rosenberg, O.S., 2017. Chlamydia interfere with an interaction between the mannose-6-phosphate receptor and sorting nexins to counteract host restriction. eLife 6, e22709. 10.7554/elife.22709

Finelli, M.J., Oliver, P.L., 2017. TLDc proteins: new players in the oxidative stress response and neurological disease. Mamm Genome 28, 395–406. 10.1007/s00335-017-9706-7

Frost, A., Unger, V.M., Camilli, P.D., 2009. The BAR domain superfamily: membrane-molding macromolecules. Cell 137, 191–196. 10.1016/j.cell.2009.04.010

Gillooly, D.J., Raiborg, C., Stenmark, H., 2003. Phosphatidylinositol 3-phosphate is found in microdomains of early endosomes. Histochem Cell Biol 120, 445–453. 10.1007/s00418-003-0591-7

Helenius, A., 1992. Unpacking the incoming influenza virus. Cell 69, 577–578. 10.1016/0092-8674(92)90219-3

Kabsch, W., 2010. XDS. Acta Crystallogr D Biol Crystallogr 66, 125–132. 10.1107/s0907444909047337

Khan, H., Winstone, H., Jimenez-Guardeño, J.M., Graham, C., Doores, K.J., Goujon, C., Matthews, D.A., Davidson, A.D., Rihn, S.J., Palmarini, M., Neil, S.J.D., Malim, M.H., 2021. TMPRSS2 promotes SARS-CoV-2 evasion from NCOA7-mediated restriction. PLoS Pathog 17, e1009820. 10.1371/journal.ppat.1009820

Khan, Md.M., Lee, S., Couoh-Cardel, S., Oot, R.A., Kim, H., Wilkens, S., Roh, S., 2022. Oxidative stress protein Oxr1 promotes V-ATPase holoenzyme disassembly in catalytic activity-independent manner. The EMBO Journal 41, e109360. 10.15252/embj.2021109360

Kvainickas, A., Jimenez-Orgaz, A., Nägele, H., Hu, Z., Dengjel, J., Steinberg, F., 2017. Cargo-selective SNX-BAR proteins mediate retromer trimer independent retrograde transport. J Cell Biol 216, 3677–3693. 10.1083/jcb.201702137

Liebschner, D., Afonine, P.V., Baker, M.L., Bunkóczi, G., Chen, V.B., Croll, T.I., Hintze, B., Hung, L.-W., Jain, S., McCoy, A.J., Moriarty, N.W., Oeffner, R.D., Poon, B.K., Prisant, M.G., Read, R.J., Richardson, J.S., Richardson, D.C., Sammito, M.D., Sobolev, O.V., Stockwell, D.H., Terwilliger, T.C., Urzhumtsev, A.G., Videau, L.L., Williams, C.J., Adams, P.D., 2019. Macromolecular structure determination using X-rays, neutrons and electrons: recent developments in *Phenix*. Acta Crystallogr D Struct Biol 75, 861–877. 10.1107/s2059798319011471

Matlin, K.S., Reggio, H., Helenius, A., Simons, K., 1981. Infectious entry pathway of influenza virus in a canine kidney cell line. J Cell Biol 91, 601–613. 10.1083/jcb.91.3.601

Maxson, M.E., Grinstein, S., 2014. The vacuolar-type H+-ATPase at a glance - more than a proton pump. Journal of Cell Science 127, 4987–4993. 10.1242/jcs.158550

McCoy, A.J., 2007. Solving structures of protein complexes by molecular replacement with *Phaser*. Acta Crystallogr D Biol Crystallogr 63, 32–41. 10.1107/s0907444906045975

McKellar, J., Rebendenne, A., Wencker, M., Moncorgé, O., Goujon, C., 2021. Mammalian and Avian Host Cell Influenza A Restriction Factors. Viruses 13, 522. 10.3390/v13030522

Merkulova, M., Păunescu, T.G., Azroyan, A., Marshansky, V., Breton, S., Brown, D., 2015. Mapping the H+ (V)-ATPase interactome: identification of proteins involved in trafficking, folding, assembly and phosphorylation. Sci Rep 5, 14827. 10.1038/srep14827

Mirrashidi, K.M., Elwell, C.A., Verschueren, E., Johnson, J.R., Frando, A., Von Dollen, J., Rosenberg, O., Gulbahce, N., Jang, G., Johnson, T., Jäger, S., Gopalakrishnan, A.M., Sherry, J., Dunn, J.D., Olive, A., Penn, B., Shales, M., Cox, J.S., Starnbach, M.N., Derre, I., Valdivia, R., Krogan, N.J., Engel, J., 2015. Global Mapping of the Inc-Human Interactome Reveals that Retromer Restricts Chlamydia Infection. Cell Host & Microbe 18, 109–121. 10.1016/j.chom.2015.06.004

Oot, R.A., Wilkens, S., 2024. Human V-ATPase function is positively and negatively regulated by TLDc proteins. Structure 32, 989–1000.e6. 10.1016/j.str.2024.03.009

Paul, B., Kim, H.S., Kerr, M.C., Huston, W.M., Teasdale, R.D., Collins, B.M., 2017. Structural basis for the hijacking of endosomal sorting nexin proteins by Chlamydia trachomatis. eLife 6, e22311. 10.7554/elife.22311

Pinto, L.H., Holsinger, L.J., Lamb, R.A., 1992. Influenza virus M2 protein has ion channel activity. Cell 69, 517–528. 10.1016/0092-8674(92)90452-i

Pinto, L.H., Lamb, R.A., 2006. The M2 proton channels of influenza A and B viruses. J Biol Chem 281, 8997–9000. 10.1074/jbc.r500020200

Simonetti, B., Danson, C.M., Heesom, K.J., Cullen, P.J., 2017. Sequence-dependent cargo recognition by SNX-BARs mediates retromer-independent transport of CI-MPR. J Cell Biol 216, 3695–3712. 10.1083/jcb.201703015

Simonetti, B., Paul, B., Chaudhari, K., Weeratunga, S., Steinberg, F., Gorla, M., Heesom, K.J., Bashaw, G.J., Collins, B.M., Cullen, P.J., 2019. Molecular identification of a BAR domain-containing coat complex for endosomal recycling of transmembrane proteins. Nat Cell Biol 21, 1219–1233. 10.1038/s41556-019-0393-3

Sirvent, A., Vigy, O., Orsetti, B., Urbach, S., Roche, S., 2012. Analysis of SRC oncogenic signaling in colorectal cancer by stable isotope labeling with heavy amino acids in mouse xenografts. Mol Cell Proteomics 11, 1937–1950. 10.1074/mcp.m112.018168

Stauffer, S., Feng, Y., Nebioglu, F., Heilig, R., Picotti, P., Helenius, A., 2014. Stepwise priming by acidic pH and a high K+ concentration is required for efficient uncoating of influenza A virus cores after penetration. J Virol 88, 13029–13046. 10.1128/jvi.01430-14

Sun, Q., Yong, X., Sun, X., Yang, F., Dai, Z., Gong, Y., Zhou, L., Zhang, X., Niu, D., Dai, L., Liu, J.-J., Jia, D., 2017. Structural and functional insights into sorting nexin 5/6 interaction with bacterial effector IncE. Sig Transduct Target Ther 2, 17030. 10.1038/sigtrans.2017.30

Villalón-Letelier, F., Brooks, A.G., Saunders, P.M., Londrigan, S.L., Reading, P.C., 2017. Host Cell Restriction Factors that Limit Influenza A Infection. Viruses 9, 376. 10.3390/v9120376

Wang, R., Qin, Y., Xie, X.-S., Li, X., 2022. Molecular basis of mEAK7-mediated human V-ATPase regulation. Nat Commun 13, 3272. 10.1038/s41467-022-30899-z

Wassmer, T., Attar, N., Bujny, M.V., Oakley, J., Traer, C.J., Cullen, P.J., 2006. A loss-of-function screen reveals SNX5 and SNX6 as potential components of the mammalian retromer. Journal of Cell Science 120, 45–54. 10.1242/jcs.03302

Weering, J.R.T. van, Verkade, P., Cullen, P.J., 2010. SNX-BAR proteins in phosphoinositide-mediated, tubular-based endosomal sorting. Semin Cell Dev Biol 21, 371–380. 10.1016/j.semcdb.2009.11.009

Wharton, S.A., Belshe, R.B., Skehel, J.J., Hay, A.J., 1994. Role of virion M2 protein in influenza virus uncoating: specific reduction in the rate of membrane fusion between virus and liposomes by amantadine. J Gen Virol 75 ( Pt 4), 945–948. 10.1099/0022-1317-75-4-945

Wickenhagen, A., Sugrue, E., Lytras, S., Kuchi, S., Noerenberg, M., Turnbull, M.L., Loney, C., Herder, V., Allan, J., Jarmson, I., Cameron-Ruiz, N., Varjak, M., Pinto, R.M., Lee, J.Y., Iselin, L., Palmalux, N., Stewart, D.G., Swingler, S., Greenwood, E.J.D., Crozier, T.W.M., Gu, Q., Davies, E.L., Clohisey, S., Wang, B., Costa, F.T.M., Santana, M.F., Ferreira, L.C.D.L., Murphy, L., Fawkes, A., Meynert, A., Grimes, G., Investigators, ISARIC4C, Filho, J.L.D.S., Marti, M., Hughes, J., Stanton, R.J., Wang, E.C.Y., Ho, A., Davis, I., Jarrett, R.F., Castello, A., Robertson, D.L., Semple, M.G., Openshaw, P.J.M., Palmarini, M., Lehner, P.J., Baillie, J.K., Rihn, S.J., Wilson, S.J., 2021. A prenylated dsRNA sensor protects against severe COVID-19. Science 374, eabj3624. 10.1126/science.abj3624

Yong, X., Zhao, L., Deng, W., Sun, H., Zhou, X., Mao, L., Hu, W., Shen, X., Sun, Q., Billadeau, D.D., Xue, Y., Jia, D., 2020. Mechanism of cargo recognition by retromer-linked SNX-BAR proteins. PLoS Biol 18, e3000631. 10.1371/journal.pbio.3000631

Yu, J.W., Lemmon, M.A., 2001. All Phox Homology (PX) Domains from Saccharomyces cerevisiae Specifically Recognize Phosphatidylinositol 3-Phosphate. Journal of Biological Chemistry 276, 44179–44184. 10.1074/jbc.m108811200

Yu, L., Croze, E., Yamaguchi, K.D., Tran, T., Reder, A.T., Litvak, V., Volkert, M.R., 2015. Induction of a Unique Isoform of the *NCOA7* Oxidation Resistance Gene by Interferon β-1b. Journal of Interferon & Cytokine Research 35, 186–199. 10.1089/jir.2014.0115

Zhirnov, O.P., 1990. Solubilization of matrix protein M1/M from virions occurs at different pH for orthomyxo- and paramyxoviruses. Virology 176, 274–279. 10.1016/0042-6822(90)90253-n

